# Investigating the mechanistic role of painful self-experience in emotional contagion: an effect of auto-conditioning?

**DOI:** 10.1101/2022.06.27.497737

**Authors:** Julian Packheiser, Efe Soyman, Enrica Paradiso, Eline Ramaaker, Neslihan Sahin, Sharmistha Muralidharan, Markus Wöhr, Valeria Gazzola, Christian Keysers

## Abstract

Emotional contagion refers to the transmission of emotions from one conspecific to another. Previous research in rodents has demonstrated that the self-experience of footshocks enhances how much an observer is affected by the emotional state of a conspecific in pain or distress. We hypothesized auditory auto-conditioning to contribute to this enhancement: during the observer’s own experience of shocks, the animal associates its own audible nocifensive responses, i.e. its pain squeaks, with the negative affective state induced by the shock. When the animal later witnesses a cage mate receive shocks and hears it squeak, the previously strengthened connection between fear and squeaks could be a mechanism eliciting the enhanced fearful response in the observer. As hypothesized, in a first study, we found pre-exposure to shocks to increase freezing and 22 kHz vocalizations associated with distress upon the playback of pain squeaks. Freezing was also increased during the playbacks of phase-scrambled squeaks, but 22 kHz calls were more frequent during the playback of regular squeaks. Core to the notion of auto-conditioning is that the effect of pre-exposure is due to the pairing of a pain-state with hearing one’s own pain squeaks. In a second study, we therefore compared the response to squeak playbacks after animals had been pre-exposed to pairings of a CO2 laser with a squeak playback against three control groups that were pre-exposed to the CO2 laser alone, to squeak playbacks alone or to neither of these conditions. We however could not find any differences in freezing or 22 kHz calls among all experimental groups. In summary, we demonstrate the sufficiency of pain squeaks to trigger fear in a way that critically depends on the nature of an animal’s prior experience and discuss why the pairing of a CO2 laser with pain squeaks cannot substitute footshock pre-exposure.

## Introduction

Empathy refers to the ability to understand and share the feelings of other individuals (Baron-Cohen & Wheelwright, 2004; Keysers & Gazzola, 2014a; Paradiso et al., 2021). The *understanding* comprises the cognitive component of empathy whereas the *sharing* comprises the affective component of empathy, which, in its simplest form, is also known as emotional contagion. Emotional contagion describes the tendency for emotional states to transmit from one individual to another leading to a convergence of emotional states (Decety & Ickes, 2011; Hatfield et al., 1993; Keysers et al., 2022). Emotional contagion has been documented to be widespread in the animal kingdom and is considered a pre-cursor to empathy (Pérez-Manrique & Gomila, 2022).

Research on emotional contagion in non-human animals has largely focused on emotional contagion of negative affective states such as pain or fear (Keysers et al., 2022). In both mice and rats, several studies have demonstrated that behavior indicative of fearful states, such as freezing, can be triggered by witnessing a demonstrator in distress (Allsop et al., 2018; Atsak et al., 2011; Carrillo et al., 2015; Jeon et al., 2010; Keum et al., 2018; Kim et al., 2010; Terranova et al., 2022; Wöhr & Schwarting, 2008). If the demonstrator is exposed to numerous strong shocks, even observers that have not been pre-exposed to shocks show some level of freezing while witnessing the demonstrator receive shocks (Han et al., 2019; Jeon et al., 2010; Keum et al., 2018; Wöhr & Schwarting, 2008). However, the response is strengthened if the observer has previously been pre-exposed to footshocks (Atsak et al., 2011; Han et al., 2019; Terranova et al., 2022). Interestingly, while in naïve observers the transmission of fear appears to depend strongly on vision (Jeon et al., 2010; Terranova et al., 2022), the pre-exposure appears to make animals more sensitive to auditory cues (Kim et al., 2010; Terranova et al., 2022).

How the prior experience of footshocks can sensitize animals to respond to the distress of others remains incompletely understood. One mechanistic proposal is based on Hebbian learning and auto-conditioning (Cruz et al., 2020; Keysers et al., 2014; Keysers & Gazzola, 2014b; Kim et al., 2010; Parsana, Moran, et al., 2012). In this framework, it is hypothesized that animals receiving an aversive stimulus react to it with nocifensive responses such as freezing, jumping and emitting vocalizations in the ultrasonic (around 22kHz) and audible range (pain squeaks). During this experience, the animals could strengthen synaptic connections between the neural populations associated with the aversive inner state and nocifensive reactions with populations encoding the sensory input triggered by witnessing their own responses – including the silence triggered by freezing, and the sounds of jumping and vocalizing. If the so pre-exposed animal later witnesses another animal emit similar sensory cues, the neural associations can then trigger matching internal states and nocifensive responses. The pairing of sensory cues and aversive inner states is then a form of conditioning called auto-conditioning. Evidence that rodents may auto-condition to their own freezing responses comes from a study in which naïve and shock pre-exposed rats were exposed to playbacks of movement sounds of other rats (Pereira et al., 2012). Upon listening to the silence intervals indicative of freezing by another animal, only rats with prior experience showed freezing. These results were complemented by a follow-up study in which the specific contributions of the rat’
ss own freezing during pre-exposure on emotional contagion were investigated (Cruz et al., 2020). In this study, rats underwent different methods of shock pre-exposure prior to an emotional contagion test. In one condition, the animals received shocks at the end of the pre-exposure session and were immediately taken out of the experimental chamber resulting in an aversive experience without association of freezing with the aversive stimulus. In another condition, rats received spaced shocks leading to a pronounced freezing response during intershock intervals. In the emotional contagion test, only rats that had experienced the shock event in combination with their own freezing responses showed fear contagion indicating that rats appeared to have auto-conditioned by associating the sound of their own freezing to their fear state and matching freezing response.

Auto-conditioning has also been investigated in the context of another behavioral expression of fear. When in distress due to the presence of a predator or an encounter of an aversive stimulus (Brudzynski et al., 1993), rats emit distinctive calls in the 22 kHz frequency range that usually accompany freezing behavior (Choi & Brown, 2003). Interestingly, naïve rats do not seem to respond with fear responses when hearing these calls (Parsana, Li, et al., 2012), but quickly associate them with aversive stimuli (Endres et al., 2007). Lesioning the auditory pathway prior to shock pre-exposure and a subsequent emotional contagion test leads to decreased rates of freezing indicating that auto-conditioning to the 22 kHz calls could also mediate the transmission of fear for this behavior (Kim et al., 2010). The auto-conditioning hypothesis to these calls was tested by (Parsana, Li, et al., 2012) who played back either calls in the 22 kHz range or rat “laughter” in the 50 kHz range to animals with or without previous self-experience with footshocks. Only rats pre-exposed to shock showed fearful responses that were selective for stimuli in the 22 kHz range, supporting the auto-conditioning hypothesis. It should be noted however that a follow-up experiment failed to support this hypothesis as freezing rates during 22 kHz call playback were identical between animals that were vocalizing or non-vocalizing in the pre-exposure session (Calub et al., 2018).

A third self-produced signal that the animals could potentially auto-condition to are pain squeaks. In rats, these squeaks are invariably emitted when animals are expose to stronger footshocks as a signal of pain (Jourdan et al., 1995) and are thus another viable stimulus that could mediate the effect of pre-exposure on emotional contagion. In the present study, we aimed to test the auto-conditioning hypothesis for pain squeaks. We first investigated the role of shock pre-exposure on fear responses during an auditory squeak playback test. We hypothesized that rats auto-condition to their own squeaking during the pre-exposure, and thus elicit more freezing responses and 22 kHz calls upon hearing the squeaks of other animals. We also predicted that this response is specific to squeaking and does not occur when control auditory stimuli are played to rats with shock pre-exposure.

## Methods Experiment 1

### Subjects

Twenty-four adult male Long Evans rats (6 – 8 weeks old; 250 – 350g; Janvier, France) were used as experimental subjects in the first experiment. The animals were randomly assigned to the experimental groups upon arrival in the local animal facility at the Netherlands Institute for Neuroscience where animals were housed socially (Type IV cages with two to four animals per cage) with *ad libitum* access to food and water in a specific pathogen free room controlled for temperature (22 – 24 °C), relative humidity (55%), and lighting (12h reversed light/dark cycle). All experimental procedures were approved by the Centrale Commissie Dierproeven (CCD number: AVD801002015105) and by the welfare body at the Netherlands Institute for Neuroscience (study dossier number: NIN181109).

### Experimental Groups

Animals were divided into three experimental groups based on the pre-exposure condition and the test stimuli to which they were subjected. Animals in two of these groups were pre-exposed to footshocks prior to the auditory playback tests. During the tests, one of these groups was presented with previously recorded squeak vocalizations (Shock->Squeak group, n=10), whereas the other group was presented with control stimuli synthesized from the original squeaks (Shock->Control group, n=9, see *Stimuli* for details). Animals in the third group were not pre-exposed to footshocks and were tested with the original squeaks (NoShock->Squeak group, n = 5).

### Stimuli

Original squeak vocalizations were recorded from adult male Long-Evans rats receiving footshocks (1s, 1.5mA) during an emotional contagion test published elsewhere (Han et al., 2019) using a CM16/CMPA condenser ultrasound microphone with a UltraSoundGate 116Hn audio recording system and the Avisoft-RECORDER software (Avisoft Bioacoustics, Germany). Five different squeak exemplars were recorded from each of three different rats for generalizability. These recordings were manually trimmed, tapered with a Tukey window, and root-mean-square-amplitude-normalized over the entire duration, resulting in 15 individual squeaks with a 1.077s mean duration (SD = ±0.096s). The control stimuli were synthesized from the original squeaks via Fourier-transforming the signal first, then randomly shuffling the phase spectrum, and finally inverse Fourier-transforming the signal. This procedure ensures that the temporal structure of the sound is entirely taken out, while the spectral structure over the whole duration of the sound remains intact. We chose to use these phase-scrambled squeaks as control stimuli because our pilot studies suggested that their playback elicited reduced levels of freezing. Any experimental rat was only presented with the original or the phase-scrambled versions of the five squeaks of only one of the three rats from which the squeaks were recorded. An example squeak, together with the corresponding phase-scrambled control version, is shown in Figure 1.

**Figure 1.**
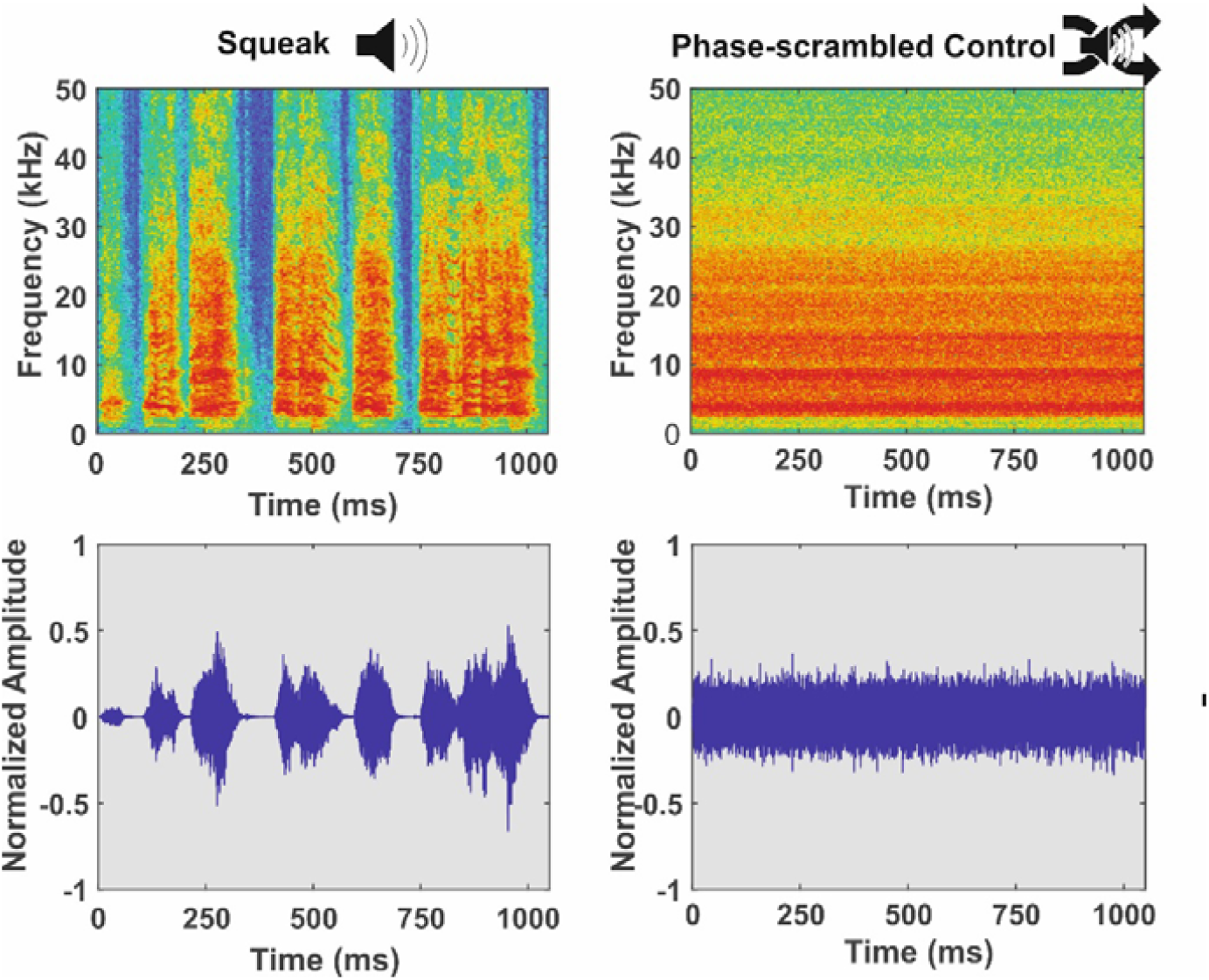
Employed stimulus material. Top row: Spectrogram of a regular squeak (left panel) and a phase-scrambled squeak (right panel). Bottom row: Normalized amplitude of a regular squeak (left panel) and a phase-scrambled squeak (right panel).

The sound pressure levels of the original squeaks were measured to be 90 dB on average (Range: 88-92 dB) by a microphone located above the center point of the observer chamber in the emotional contagion setup described in Han et al. (2019). These sound pressure levels were quantified using the Calibrated 40 kHz Reference Signal Generator in combination with the Avisoft-SASLabPro software version 5.2.13 (Avisoft Bioacoustics, Germany). Specifically, the sound pressure level of each squeak was measured by fırst automatically segmenting the individual bouts that make up the squeak, then calculating the root-mean-square-amplitude of each bout in dB, and finally taking the average dB of all the bouts. In the current study, all stimuli were played back at either 75 or 85 dB. This difference was initially used to investigate possible effects of amplitude on animals’ fear responses. In total, five animals from the Shock->Squeak and the Shock->Control group and no animal from the NoShock->Squeak group received squeak playbacks at 75 dB. Five animals from the Shock->Squeak group, four animals from the Shock->Control group and five animals from the Naive->Squeak group were presented with stimuli at 85 dB (see Table 1 for a summary). We decided to pool the results of these two amplitude levels as they led to highly comparable patterns of fear responses. To account for this experimental difference, we included the amplitude of the playback as a covariate in the statistical model (see *Statistical Analysis for details*).

**Table 1.** Number of animals per group with respect to different amplitudes during playback.

|  | Playback at 75 dB | Playback at 85 dB |
| --- | --- | --- |
| **Shock -> Squeak** | n = 5 | n = 5 |
| **Shock -> Control** | n = 5 | n = 4 |
| **Naïve -> Shock** | n = 0 | n = 5 |

### Apparatus

Pre-exposure with footshocks was delivered in a custom-built pre-exposure chamber (L: 30 cm x W: 20cm x H: 40cm) featuring two experimental chambers divided by a transparent perforated separator. As contextual markers, the walls of the pre-exposure chamber were covered with black and white stripes, the overhead daylights were turned on, the background radio was turned off, and the chamber was wiped with a vanilla aroma after cleaning with a rose-scented dishwashing soap. During the experimental procedure, the animals were placed on stainless steel grid rods of one of the experimental chambers through which electrical currents could be applied to the animals via a stimulus scrambler (ENV 414-S, Med Associates Inc., VT). Auditory playbacks were administered in another room in a different test chamber (L: 24 cmx W: 25cm x H: 34cm) consisting of two adjacent compartments with a transparent perforated divider in between. This test context differed from the pre-exposure context to avoid contextual fear conditioning: the walls of the testing chamber were made of transparent Plexiglass, the lights were turned off, the background radio was turned on at low levels, and a lemon-scented dishwashing soap was applied to the chamber after cleaning with 70% ethanol. Behavior was recorded using a Basler GigE camera (acA 1300-60g), which was mounted to the ceiling of the test chamber and controlled by EthoVisionXT (Noldus, the Netherlands). The ultrasound microphone described above was positioned on top of the compartment that the animal was in, while a Vifa ultrasonic dynamic speaker (Avisoft Bioacoustics, Germany) was positioned in the adjacent compartment facing the animal’s compartment. During both the habituation and the auditory playback test phases, bedding material from an unknown, unstressed male rat was placed in the adjacent chamber to prime the rats to the possibility that there was another rat in the vicinity.

### Experimental Procedure

After acclimatization to the local animal facilities for at least one week, the animals were handled for five minutes each day for three days. Then the experimental procedure started (Figure 2). On Day 1, all rats were habituated to the auditory playback test context by allowing them to freely explore the chamber for 20 minutes in the dark. At the end of habituation, the animals were taken out of the chamber and handled for five minutes. On Day 2, the animals underwent the shock pre-exposure procedure in the pre-exposure context in light condition. The Shock->Squeak and the Shock->Control groups received four unpredictable shocks (1s, 0.8mA) with an intershock interval of 240-360 seconds. The animals in the NoShock->Squeak group were also placed in the same chamber for the same amount of time but did not receive any shocks. At the end of pre-exposure, all animals were first handled for five minutes and then rested individually for one hour in a transportation cage to prevent negative emotional contagion in the home cage. After the pre-exposure day, there were two more habituation days in the playback test context (Day 3 and 4 in Figure 2) administered exactly as in Day 2 to reduce baseline freezing and fear in the later test session.

**Figure 2.**
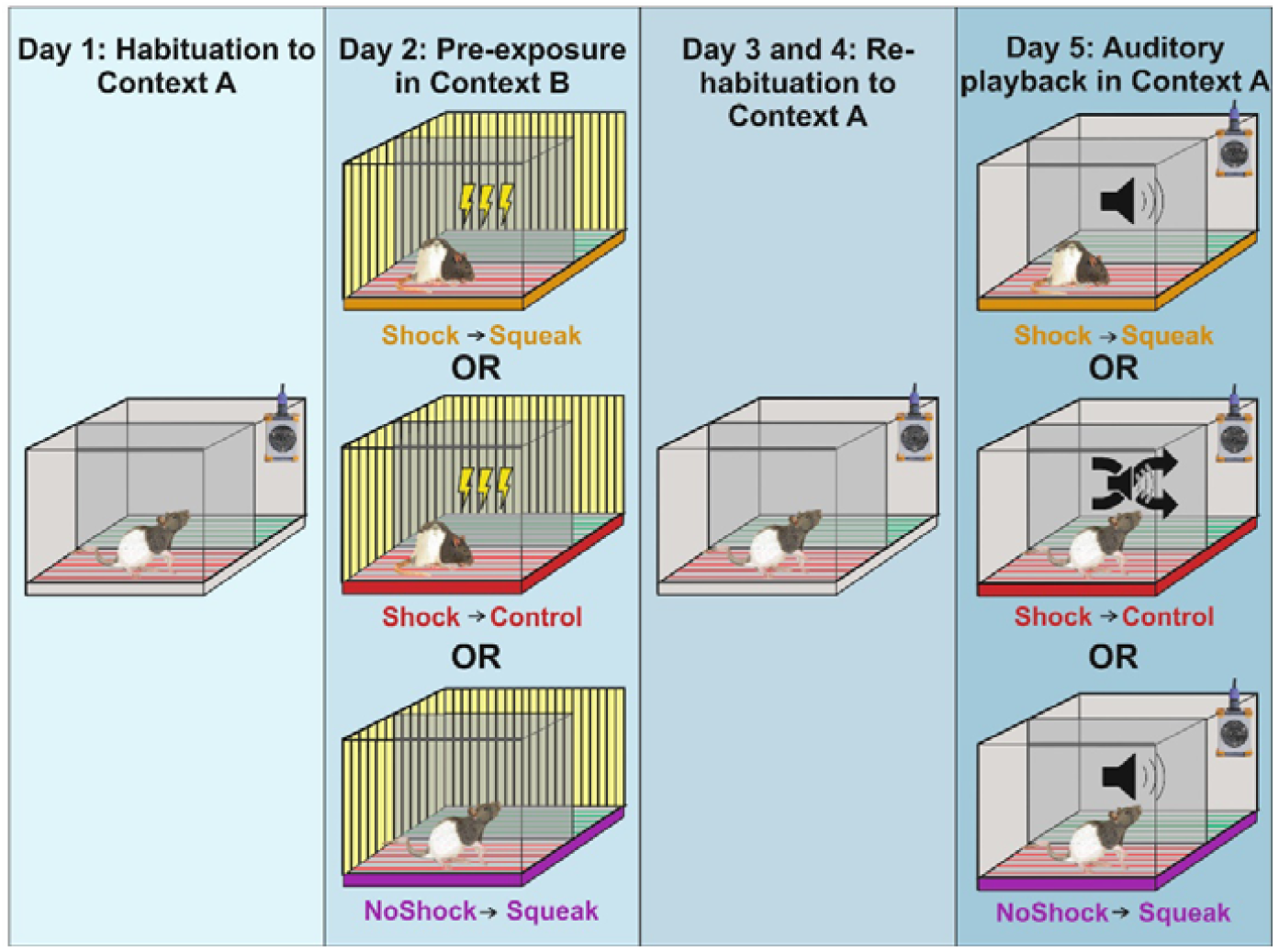
Experimental paradigm. Experiments took place in two-compartment chamber separated by a transparent divider. Pre-exposure on day 2 could either consist of footshocks (Shock->Squeak and Shock->Control groups) or a resting period (NoShock->Squeak group). Playback on day 5 was of regular squeaks (Shock->Squeak and NoShock->Squeak groups) or phase-scrambled squeaks (Shock->Control). Note that the yellow background in the pre-exposure box symbolizes the fact that overhead lights were turned on in context B but not A.

On Day 5 the auditory playback tests were administered. After a 12-minute baseline, five playbacks of either the original squeaks or the phase-scrambled control stimuli were administered with an interstimulus interval of 120-180 seconds. The timing and number of playbacks was chosen to match that of our experiments involving shocks to demonstrator in the neighboring compartment (Han et al. 2020; 2019), to enable comparison of freezing levels across experiments. Animals stayed in the test chamber for another 120 seconds after the presentation of the last stimuli. The stimuli were played back using the playback system described above with the Avisoft-RECORDER software at a sampling rate of 250 kHz. The sound amplitude levels were calibrated and adjusted with the Calibrated 40 kHz Reference Signal Generator in combination with the Avisoft-SASLabPro software version 5.2.13. At the end of the auditory playback tests, animals were housed individually for one hour to prevent stress contagion in the home cage.

### Freezing and 22 kHz Call Quantification

Freezing was scored in BORIS v 7.7.3 (Friard & Gamba, 2016) by an experimenter blind to the experimental conditions. During the auditory playback tests, an infrared LED was attached to the setup to provide visual feedback in the video recordings when a stimulus was presented. A threshold of minimally 3 seconds of freezing was used to ensure that the scored behavior was actual freezing and not just transient immobility. Percentages of time spent freezing within the baseline and the auditory playback periods (12 minutes each) were calculated separately for statistical analyses. 22 kHz vocalizations were semi-automatically detected using the MATLAB toolbox DeepSqueak (version 2.5.0, Long Rat Call_V2 network with default settings, (Coffey et al., 2019)) first, and then manually checked by an operator blind to the experimental conditions. Identically to the freezing scores, percentages of time spent emitting 22 kHz calls within the baseline and the auditory playback periods were calculated separately for statistical analyses. Opposed to experiments involving a real demonstrator where attributing vocalizations to the observer or demonstrator can be difficult, all vocalizations in our experiment (except those involved in the playbacks) could unambiguously be attributed to the observer.

### Statistical Analysis

Statistical analyses were performed using R (version 4.1.3) and JASP (version 0.16.1). In a first step, frequentist statistics were applied. Increases from baseline to playback period for each experimental group were tested by applying either paired t-tests or Wilcoxon signed-rank tests depending on normality violations measured via a Shapiro-Wilk (abbreviated as S-W) test. To determine differences between the groups, the baseline and playback period were analyzed separately using a one-way ANOVA (levels: Shock->Squeak, Shock->Control, NoShock->Squeak) if normality assumptions for the residuals were met. For the playback session, the amplitude of the squeak (75 vs 85 dB) was included as a covariate to account for the potential influence of the loudness of the auditory playback. If normality was violated (S-W p-value < .05), we instead calculated a non-parametric Kruskal-Wallis test to identify differences between the experimental groups. In an additional analysis, we also calculated either a one-way ANOVA or a Kruskal-Wallis test for the difference score between the baseline and auditory playback period to control for potential variability in baseline fear responses. This is similar to assessing the interaction effect of Epoch (Baseline vs. Playback) x Group (Shock->Squeak, Shock->Control, NoShock->Squeak), but because there is no well-established non-parametric test to examine such interactions, assessing the effect of group on the Playback-Baseline measures seemed a more robust alternative. Again, amplitude was included in the model if parametric tests could be applied. Significant main effects were investigated post hoc using parametric t-tests or non-parametric Wilcoxon rank-sum tests. All tests were conducted two-tailed. Because only two of the potential three pairwise comparisons have meaning, we used planned comparisons to assess the effect of pre-exposure (Shock->Squeak vs. NoShock->Squeak) or the specificity for squeaks (Shock->Squeak vs. Shock->Control) that do not require correction for multiple comparisons (Keppel & Zedeck, 1989).

The same factorial designs were also analyzed with Bayesian ANOVAs, which have an advantage over frequentist statistics as they can quantify not only the evidence for the presence of an effect, but also the evidence for the absence of an effect, as well as the absence of evidence for either (Keysers et al., 2020). For main effects and interactions, the BF_incl_ was used as a marker of evidence. For post hoc tests of main effects, the BF_10_ was used. For both the BF_incl_ and the BF_10_, a value of greater than 3 is considered to provide evidence in favor of the alternative hypothesis, whereas a value of less than 1/3 provides evidence in favor of the null hypothesis. Interpretation of the Bayes factors followed the guidelines by (Lee & Wagenmakers, 2014). All analyses were performed with default prior settings in JASP and effects were estimated across all models.

## Results Experiment 1

### Pre-exposure session

All animals in the Shock->Squeak and Shock->Control groups (19/19) emitted audible squeak vocalizations in response to each of the four footshocks, whereas none of the animals in the NoShock->Squeak group (0/5) emitted any squeaks during their sham pre-exposure session.

### Playback session

In a first step, we aimed to verify that fear responses were comparable between the experimental groups during the baseline period of the playback session. The experimental groups did not show differences in freezing responses (S-W p-value = 0.128; F_(2,20)_ = 2.46 *p* = 0.110, η^2^ = 0.19) or in 22 kHz call emissions (S-W p-value < 0.001; χ^2^_(2)_ = 1.31, *p* = 0.519, η^2^ = 0.03). These results were complemented by corresponding Bayesian ANOVAs which indicated either absence of evidence of an effect for freezing responses (BF_incl_ = 0.69) or moderate evidence of absence for the 22 kHz vocalizations indicative of distress (BF_incl_ = 0.30).

Next, we aimed to determine differences in fear behavior during the playback period across groups. We found significant main effects on both freezing rates (S-W p-value = 0.608; F_(2,20)_ = 4.32, *p* = 0.028, η^2^ = 0.30, BF_incl_ = 2.11) as well as 22 kHz call emissions (S-W p-value = 0.221; F_(2,20)_ = 8.39, *p* = 0.002, η^2^ = 0.45, BF_incl_ = 9.67). Planned comparisons for freezing rates showed absence of evidence for a difference between the Shock->Squeak and the Shock->Control group (t = 0.50, *p* = 0.601, BF_10_ = 0.64). There was approaching moderate evidence that freezing was increased in the Shock->Squeak compared to the NoShock->Squeak group (t = 2.56, *p* = 0.025, BF_10_ = 2.85). For 22 kHz calls, we found approaching moderate evidence for the Shock->Squeak group emitting more 22 kHz calls compared to the Shock->Control group (t = 2.41, *p* = 0.014, BF_10_ = 2.55) and moderate evidence to emit more 22 kHz calls compared to the NoShock->Squeak group (t = 3.16, *p* = 0.004, BF_10_ = 6.26). Playback amplitude neither reached significance for freezing (F_(1,20)_ = 0.10, *p* = 0.752) nor 22 kHz calls (F_(1,20)_ = 0.47, *p* = 0.500) suggesting that it did not play a role in modifying the fear responses, a result contrasting a previous study on locomotor inhibition (Fendt et al., 2018). Results for each individual experimental group are depicted in Figure 3A for freezing and in Figure 3B for 22 kHz calls. Descriptive statistics for freezing responses and 22 kHz calls are presented in Supplementary Tables 1 and 2, respectively.

**Figure 3.**
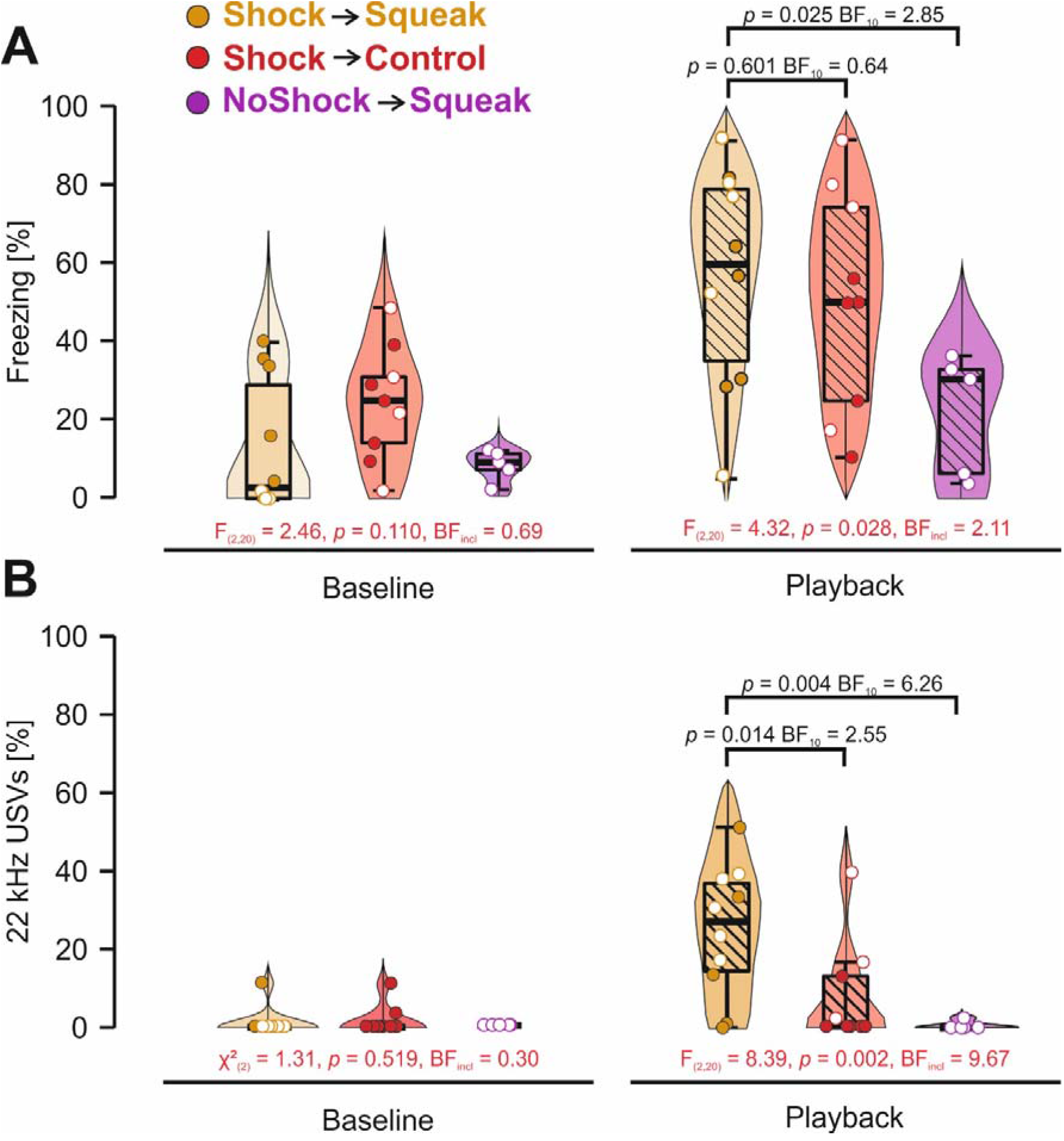
Behavioral results for the playback session on day 5. A) Proportion of freezing responses in percent during baseline and auditory playback in the playback session. Animals with higher amplitude playback are marked by an open circle. Only between group differences are presented here. Within group differences from baseline to playback are depicted in Supplementary Figure 1. **B)** Proportion of 22 kHz calls in percent during baseline and auditory playback in the playback session. Animals with higher amplitude playback are marked by an open circle. Only between group differences are presented here. Within group differences from baseline to playback are depicted in Supplementary Figure 2. * represents p < .05. ** represents p < .01.

To account for inter-individual variability in baseline fear responses, we repeated the previous analyses and used the relative increase from baseline to playback as dependent variable. We again found significant main effects and moderate evidence that freezing rates (S-W p-value = 0.657, F_(2,20)_ = 6.14, *p* = 0.008, η^2^ = 0.34, BF_incl_ = 4.71) and strong evidence that 22 kHz call emissions (S-W p-value = 0.015; χ^2^_(2)_ = 9.38, *p* = 0.007, η^2^ = 0.37, BF_incl_ = 21.02) differed between the groups. As for the data from the playback period only, there was absence of evidence that the Shock->Squeak group differed in terms of freezing compared to the Shock->Control group (*t* = 1.38, *p* = 0.149, BF_10_ = 0.61). Similarly, there was absence of evidence for a difference between the Shock->Squeak compared to the NoShock->Squeak group (*t* = 1.93, *p* = 0.045, BF_10_ = 1.38). Amplitude as covariate did not reach significance for freezing rates (F_(1,20)_ = 3.81, *p* = 0.065). For 22 kHz calls, we found a significant effect and anecdotal evidence for an increase for the Squeak->Shock group compared to the NoShock->Squeak group (U = 4, *p* = 0.009, BF_10_ = 1.94). The comparison to the Shock->Control group was significant but revealed absence of evidence (U = 17.5, *p* = 0.025, BF_10_ = 1.49).

The final analysis we conducted aimed to identify whether squeak playback elicited similar freezing responses compared to a classical emotional contagion design during which the observer is paired with another conspecific. Here, we made use of pre-existing data of nine rats from a previously published study (Han et al., 2019) and compared them to the data from the Shock->Squeak group. The direct comparison between the result patterns of these two studies is possible since the experimental protocol (*pre-exposure*: session duration, number of shocks, shock amplitude and interstimulus interval || *test session*: session duration, number of shocks/squeak playbacks) of the present study was identical to the study by Han et al. (2019). 22 kHz calls were not compared between the studies as they were not recorded in Han et al. (2019). While the freezing rates were reduced on a descriptive level during squeak playback (56.69%±27.95%) compared to a classical observation (69.68%±14.57%), an independent sample t-test showed absence of evidence for or against a group difference (S-W p-values > 0.310; t_(16)_ = 1.25, *p* = 0.229, d = 0.57, BF_10_ = 0.69, Supplementary Figure 3). For a baseline-corrected measure of freezing, a similar picture emerged (S-W p-values > 0.300; *t*_*(16)*_ = 1.39, *p* = 0.182, d = 0.64, BF_10_ = 0.78).

## Discussion Experiment 1

In the first experiment, we investigated whether rats become sensitized to squeaking after having the opportunity to associate squeaks with aversive experiences during pre-exposure to painful shocks. As expected, freezing and 22 kHz calls rates increased for the shock pre-exposed animals from the baseline to the playback period, and were generally higher for the Shock->Squeak compared to the NoShock->Squeak group during the playback period. Although only as a trend, freezing responses tended to increase to the presentation of squeaks compared to baseline in the NoShock->Squeak condition (Supplementary Figure 1). Playing back the scrambled calls elicited a similar level of freezing as regular squeak playback but lower levels of 22 kHz calls. Such responses indicate that not only auto-conditioning but also sensitization to squeaks might have been at play after being exposed to aversive stimuli (Poulos et al., 2015). Altogether these results suggest auto-conditioning may not be necessary to respond to the distress of others, but could act as an enhancer of an innate disposition to react to squeaks.

To disambiguate if the underlying mechanism facilitating emotional contagion was due to sensitization or auto-conditioning to squeaks (or possibly even both), the squeak and the aversive event need to be disentangled. To this end, we substituted the footshocks during pre-exposure, which unfortunately trigger both pain and the emission of pain squeaks, with a painful experience that has been demonstrated not to elicit squeaking: shining a CO2 laser on the animal (Carrillo et al., 2019). By pairing this painful laser experience with or without a squeak playback, auto-conditioning and sensitization should be separable: If the fear responses to later squeak playback are due to auto-conditioning, freezing rates and 22 kHz call emissions should only be increased in an experimental group that received paired pain pre-exposure with squeak playback (auto-conditioning) compared to a group with pain pre-exposure but without squeak playback pairing (sensitization). If fear responses are due to sensitization alone, both groups should show equal levels of freezing and 22 kHz calls. If both mechanisms play a role as hinted at in experiment 1, both the auto-conditioning and the sensitization group should demonstrate stronger fear responses compared to controls, and the auto-conditioning group should display stronger fear responses compared to the sensitization group.

## Methods Experiment 2

### Subjects

Eighty adult male Long Evans rats (6 – 8 weeks old; 250 – 350g; Janvier, France) were used as experimental subjects in the second experiment. As for experiment 1, the animals were randomly assigned to the experimental groups upon arrival in the local animal facility. Housing conditions were identical to experiment 1. All experimental procedures were approved by the Centrale Commissie Dierproeven (CCD numbers: AVD801002015105 and AVD8010020209724) and by the welfare body at the Netherlands Institute for Neuroscience (study dossier numbers: NIN201101 and NIN203701).

### Experimental Groups

Animals were divided into four experimental groups based on the pre-exposure condition to which they were subjected (each n = 20). Animals in one of the groups were only exposed to a painful CO2 laser stimulation during pre-exposure (Laser group), whereas the animals in another group were administered the same levels of laser simultaneously with auditory playbacks of previously recorded squeaks (Laser+Squeak group). Animals in another group received only squeak playbacks (Squeak group), while the animals in the last group were neither subjected to squeak playbacks nor painful levels of the laser (Naïve group).

### Stimuli

Stimuli used in experiment 2 during pre-exposure and auditory playback tests were identical to the original squeaks and phase-scrambled control sounds described for experiment 1. For animals that were presented with squeak playbacks during pre-exposure, different squeaks were used during the playback tests. In experiment 2, all stimuli were played back at 75 dB.

### Apparatus

Pre-exposure for experiment 2 was performed in a custom-built pre-exposure chamber different from the one used in experiment 1 (L: 30 cm x W: 15cm x H: 30cm) and placed in a different room inside a faraday cage. It consisted of a rectangular dark-colored apparatus, opened along one of the long sides with a 0.5cm fence. The opened side was placed in front of an opening in the faraday cage. The CO2 laser (CL15 model:M3) was placed outside the faraday cage and the arm used for delivering the heat pulses protruded into the faraday cage, with its tip 15cm away from the observer’s box. As in experiment 1, pre-exposure took place under normal light conditions, the background radio was turned off, and the apparatus was wiped with a vanilla aroma after cleaning with a rose-scented dishwashing soap. Auditory playback tests took place in the same test chamber and room as in experiment 1. Again, contextual differences were maximized by performing the auditory playback tests in dark conditions, turning on the radio at low volume, and applying a lemon-scent before the test session. As described for experiment 1, the ultrasound microphone and the speaker were placed in the adjacent empty chamber for recording and stimulus playback, together with bedding material from an unknown male rat.

### Experimental Procedure

All animals were acclimatized to the local animal facility and handled as in experiment 1. Animals were habituated for 10 minutes to the auditory playback test context on the first experimental day and for two consecutive days after pre-exposure (see Figure 4). For the first 40 animals (each experimental group containing ten animals) tested in this experiment, no habituation to the pre-exposure context was provided. However, due to elevated freezing responses in all experimental groups during the pre-exposure session (even in the Naïve group), the remaining 40 animals received one session of habituation in the pre-exposure context one day before the pre-exposure procedure. Furthermore, the laser was applied for this subset of animals through a hole of the closed door preventing any visual contact of the animal with the experimenter. While both these differences in procedure should have minimal impact on the auditory playback session, they might have affected the fear responses during the pre-exposure session. Thus, habituation (absent for the first 40 and present for the last 40), was included as a covariate in the statistical model for pre-exposure analyses (see *Statistical Analysis* for details).

**Figure 4.**
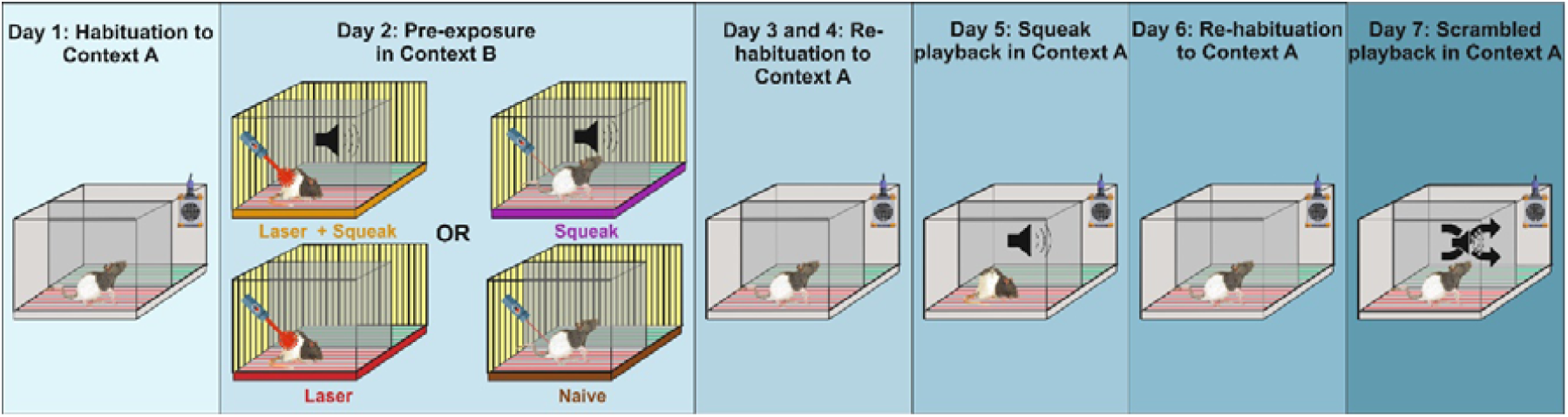
Behavioral paradigm of the second experiment. The procedure only differed between groups on the day during pre-exposure. Here, groups either received laser stimulation together with squeak playback (Laser+Squeak), only laser stimulation (Laser), only squeak playback with a low intensity laser stimulation (Squeak) or a low intensity laser stimulation only (Naïve). On day 5, regular squeaks were played back to all animals whereas phase-scrambled squeaks were played back on day 7. Note that the yellow background in the pre-exposure box represents the turned on overhead lights.

For all groups, pre-exposure started with a 2-minute baseline period, in which the animals were administered six below-threshold (30% of laser power) laser stimulations (200 msec) to habituate them to the laser arm entering the chamber and the click sound associated with laser delivery. For the Laser group, four laser stimulations at 70% intensity of the laser power to one of the paws of the animal were then administered with an intertrial interval of 240-360 seconds. This stimulation level of the C02 laser was previously shown to be effective in eliciting pain (Carrillo et al., 2019). For the Laser+Squeak group, the same procedure was applied with the addition of the playback of a previously recorded squeak that started at the same time as each laser stimulation. Here, the laser stimulation triggered squeak replay with close to zero latency to simulate a situation similar to shock-experience, in which pain squeaks are measured with short latency following the onset of footshocks (Carrillo et al., 2019). For the Squeak group, the laser intensity was reduced to the below-threshold level, which does not elicit pain (30% of laser power), and squeaks were played back as described above. Finally, for the Naive group, laser intensity was again reduced to the below-threshold levels and no squeaks were played back upon stimulation. Low intensity laser was used instead of no laser stimulation alone, to control for the aiming of the laser onto the animal and the faint clicking associated with delivering the laser. After the last stimulus delivery, the animals were left for four minutes in the apparatus before being handled for five minutes and finally were rested individually in a separate room in a transportation cage for two hours to prevent stress contagion in the home cage that they shared with two or three other rats (depending on the batch size) of the same experimental condition.

Auditory playback tests were performed exactly as described for experiment 1 except that on day 5 following the third habituation, all groups were first tested with squeak playbacks. On day 6, all animals were re-introduced to the test chamber and exposed to the context for 10 minutes to extinguish any potential negative association that might have formed due to squeak playbacks. On day 7, all groups were this time tested with phase-scrambled control sounds. Thus, in contrast to experiment 1, responses to squeaks and control sounds were tested serially in a within-subject, rather than in a between-subject design.

### Pain, Freezing and 22 kHz Call Quantification

For the second experiment, we first explored the behavioral responses during pre-exposure to validate that pain was reliably induced via laser delivery as the application of a CO2 laser as an aversive tool is less common (Kung et al., 2003; Shyu et al., 2003). We calculated a custom pain score across all four laser stimulations based on the behavioral responses of the animals following stimulus presentation. Here, animals received a score of 0 if they did not react at all, a score of 1 if they slightly twitched without retracting the limb, a score of 2 if they retracted the targeted limb and a score of 3 if they retracted the limb and moved from their current location. Thus, the total pain score could vary between 0 (minimum) and 12 (maximum). Quantification of freezing and 22 kHz calls were performed exactly as described for experiment 1, except that an additional side view camera in addition to the top view camera, was available for scoring freezing in the auditory playback tests.

### Statistical Analysis

To determine whether the experimental groups differed in terms of pain scores, freezing or 22 kHz calls during pre-exposure, a parametric one-way (Bayesian) ANOVA or a non-parametric Kruskal-Wallis test with the between-subjects factor Group (Laser+Squeak, Laser, Squeak, Naive) were calculated. Habituation to the pre-exposure context (Yes, No) was included as a covariate if normality assumptions were not violated. Video data (pain and freezing) for two animals in the Naïve, two animals in the Laser+Squeak and one animal in the Squeak condition could not be evaluated due to technical issues in the pre-exposure session.

For the auditory playback sessions using squeaks (day 5) and control sounds (day 7), we repeated the analysis procedure from experiment 1. First, increases from baseline to playback period for each experimental group were tested by applying either paired t-tests or Wilcoxon signed-rank tests. Then, we computed one-way (Bayesian) ANOVAs or Kruskal-Wallis tests with the between-subjects factor Group (Laser+Squeak, Laser, Squeak, Naive) for the baseline period using both freezing and 22 kHz calls as dependent variable. This was then repeated for the data from the playback period and for a difference score subtracting the baseline from the playback data. Analysis for the squeak sound playback (day 5) and control sound playback (day 7) was analyzed separately. Audio data from three animals (one animal from the Laser+Squeak, Laser, and Naïve groups each) from the squeak playback session and six animals (one animal from the Laser+Squeak and Naïve groups each and two animals from the Laser and Squeak groups each) for the control playback session could not be evaluated due to technical issues.

## Results Experiment 2

### Pre-exposure (Day 2)

#### Pain responses

The one-way ANOVA with the different experimental groups as factor levels revealed a significant effect (S-W p-value = 0.158; F_(3,70)_ = 134.94, *p* < 0.001, η^2^ = 0.85, BF_incl_ > 100, Figure 5A). Bayesian post hoc tests revealed extreme evidence for stronger pain reactions in both groups that received pain stimulations (Laser+Squeak and Laser) compared to the other two groups (Squeak and Naïve, all ts > 10, *p*s < 0.001, BF_10_s > 100). We found approaching moderate evidence that the laser conditions did not differ from each other (t = 0.52 = *p* = 0.603, BF_10_ = 0.35). In contrast, we found approaching moderate evidence that the two control conditions differed with respect to their pain reactions: the Squeak group showed stronger responses compared to the Naive group (t = 2.34, *p* = 0.066, BF_10_ = 2.50), possibly due to the animals moving after the squeak onset. Habituation as a covariate did not reach significance (F_(1,70)_ = 0.47, *p* = 0.356). Descriptive statistics for pain responses during pre-exposure are presented in Supplementary Table 3.

**Figure 5.**
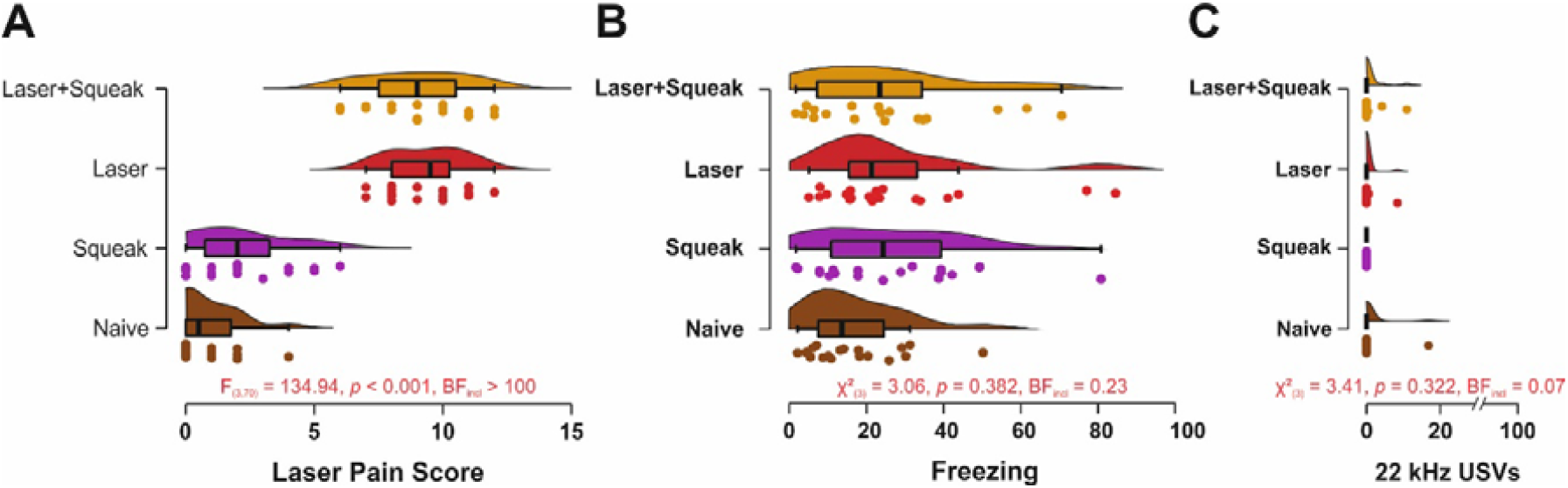
Pain and fear responses during pre-exposure on day 2. A) Cumulative pain responses to the laser for the four experimental groups. **B)** Proportion of freezing and **C)** proportion of 22 kHz calls for the different experimental groups.

#### Fear responses

Opposed to the clear pain reactions observable during high intensity laser exposure, we found moderate evidence of absence that the groups differed in freezing (S-W p-value < 0.001; χ^2^_(3)_ = 3.06, *p* = 0.382, η^2^ < 0.01, BF_incl_ = 0.23, Figure 5B) as well as strong evidence of absence that that the experimental groups emitted different levels of 22 kHz calls (S-W p-value < 0.001; χ^2^_(3)_ = 3.41, *p* = 0.322, η^2^ < 0.01, BF_incl_ = 0.07, Figure 5C). No animals emitted any squeaks over the course of pre-exposure. Descriptive statistics for freezing responses and 22 kHz call emissions during pre-exposure are presented in Supplementary Tables 4 and 5, respectively.

### Squeak Playback (Day 5)

During the squeak playback session on day 5, we again first investigated baseline differences in fear responses. Evidence against differences between the experimental groups was moderate for freezing (S-W p-value < 0.001; χ^2^_(3)_ = 1.06, *p* = 0.788, η^2^ < 0.01, BF_incl_ = 0.16) and 22 kHz calls (S-W p-value < 0.001; χ^2^_(3)_ = 4.05, *p* = 0.256, η^2^ = 0.01, BF_incl_ = 0.24). Opposed to our expectations, there was also anecdotal to moderate evidence against group differences during the playback session (*freezing*: S-W p-value < 0.001; χ^2^_(3)_ = 3.13, *p* = 0.373, η^2^ = 0.01, BF_incl_ = 0.16; *22 kHz calls*: S-W p-value < 0.001; χ^2^_(3)_ = 5.89, *p* = 0.117, η^2^ = 0.04, BF_incl_ = 0.40) as well as for the difference scores between playback and baseline (*freezing*: S-W p-value < 0.001; χ^2^_(3)_ = 3.45, *p* = 0.328, η^2^ = 0.01, BF_incl_ = 0.15; *22 kHz calls*: S-W p-value < 0.001; χ^2^_(3)_ = 6.63, *p* = 0.08, η^2^ = 0.05, BF_incl_ = 0.41). Results for each individual experimental group are depicted in Figure 6A for freezing and Figure 6B for 22 kHz calls. Descriptive statistics for freezing rates and 22 kHz calls during normal squeak playback are presented in Supplementary Tables 6 and 7, respectively.

**Figure 6.**
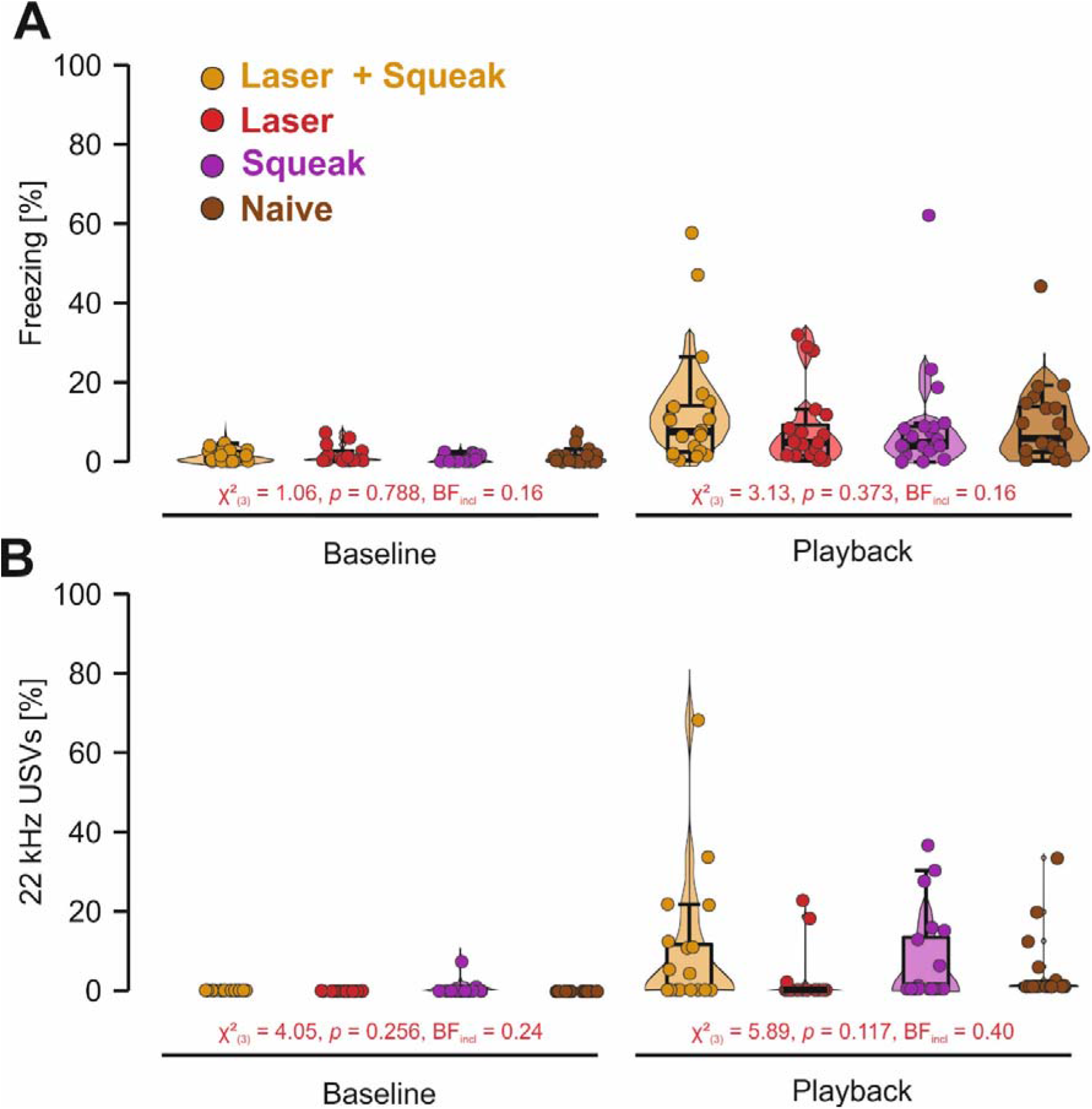
Behavioral results for the squeak playback (day 5) of experiment 2. A) Proportion of freezing responses in percent during baseline and auditory playback in the playback session on day 5. Animals with higher amplitude playback are marked by an open circle. Only between group differences are presented here. Within group differences from baseline to playback are depicted in Supplementary Figure 4. **B)** Proportion of 22 kHz vocalizations in percent during baseline and auditory playback in the playback session on day 5. Animals with higher amplitude playback are marked by an open circle. Only between group differences are presented here. Within group differences from baseline to playback are depicted in Supplementary Figure 5.

### Phase-scrambled Squeak Playback (Day 7)

During the phase-scrambled squeak playback session on day 7, we replicated the analysis procedure for the normal squeak playback. As before, we found anecdotal to strong evidence against differences between the experimental groups for freezing (S-W p-value < 0.001; χ^2^_(3)_ = 0.69, *p* = 0.788, η^2^ < 0.01, BF_incl_ = 0.13) and 22 kHz calls (S-W p-value < 0.001; χ^2^_(3)_ = 2.55, *p* = 0.466, η^2^ = 0.01, BF_incl_ = 0.70) during the baseline. Similarly, there was moderate to strong evidence against group differences during the playback session (*freezing*: S-W p-value < 0.001; χ^2^_(3)_ = 0.30, *p* = 0.961, η^2^ < 0.01, BF_incl_ = 0.09; *22 kHz calls*: S-W p-value < 0.001; χ^2^_(3)_ = 2.00, *p* = 0.573, η^2^ = 0.01, BF_incl_ = 0.11) and for the difference scores between playback and baseline (*freezing*: S-W p-value < 0.001; χ^2^_(3)_ = 0.60, *p* = 0.896, η^2^ < 0.01, BF_incl_ = 0.09; *22 kHz calls*: S-W p-value < 0.001; χ^2^_(3)_ = 3.31, *p* = 0.346, η^2^ < 0.01, BF_incl_ = 0.11). Results for each individual experimental group are depicted in Supplementary Figure 6A for freezing and Supplementary Figure 6B for 22 kHz calls. Descriptive statistics for freezing rates and 22 kHz calls during on phrase-scrambled squeak playback are presented in Supplementary Tables 8 and 9, respectively.

### Cross-experimental Comparison of Fear Responses during Squeak Playback

In a final exploratory analysis, we investigated whether freezing differed between experiment 1 and experiment 2 during playback in groups with aversive pre-exposure. To this end, we conducted a Wilcoxon rank sum test for freezing responses and 22 kHz calls during squeak playback between shock or laser pre-exposed animals from both experiments. Fear responses were pooled across both groups that received laser pre-exposure as there was evidence against group differences in the previous analysis. We found moderate evidence for increased freezing (S-W p-value < 0.001; W = 37.5, p < 0.001, BF_10_ = 5.05) and increased 22 kHz calls (S-W p-value < 0.001; W = 58.5, p < 0.001, BF_10_ = 3.16) after shock pre-exposure compared to laser pre-exposure.

## Discussion Experiment 2

In the second experiment, we aimed to disentangle the effect of the potential association created by pairing self-experience with painful events and squeak emissions on subsequent fear responses during squeak playbacks. We hypothesized that the fear responses would be increased upon playback if pain pre-exposure in combination with squeak playback were given compared to a group exposed only to a pain stimulus. If fear responses were due to sensitization, both groups would have shown equal levels of freezing and 22 kHz calls, but both would be higher than the pre-exposures not including pain. None of the hypotheses could be confirmed as neither freezing levels nor 22 kHz call emissions differed between the experimental groups that received a painful experience during pre-exposure and the controls without pain self-experience. These results were surprising given that CO2 laser stimulation has been previously used to induce fear conditioning (Kung et al., 2003; Shyu et al., 2003) and it has been shown to trigger emotional mirror neurons which are active during the observation of other rats receiving painful shocks (Carrillo et al., 2019). We discuss these findings within the scope of the general discussion. In addition, although all groups showed increased freezing upon playback compared to baseline, this did not differ between the intact and scrambled squeaks and did not reach the levels observed following pre-exposure to footshocks.

## General Discussion

The present study investigated the role that pain squeaks have in triggering fear responses (freezing and 22-kHz ultrasonic vocalizations), how the response to these squeaks depends on prior experience with footshocks and such pre-exposure to footshocks could be substituted by the pairing of a painful CO_2_ laser with the sound of pain squeaks, as Hebbian learning or autocoditioning perspectives may suggest. We focused on pain squeaks, because they had so far not been explored in emotional contagion paradigms, yet occur at a moment that coincides with responses in the cingulate cortex that are necessary for emotional contagion (Carrillo et al., 2019).

In the first experiment, we showed that listening to pain squeaks indeed triggered typical fear reactions in rats pre-exposed to footshocks, including an increase in freezing and emission of 22kHz ultrasonic vocalizations. Without pre-exposure, listening to pain squeaks only triggered low freezing rates and no 22 kHz calls. We further found that 22kHz ultrasonic vocalizations were almost exclusively emitted during intact squeak playbacks and not in response to phase-scrambled squeaks, suggesting a specificity for squeak vocalizations. Freezing behavior was indistinguishable between animals receiving squeak or phase-scrambled control stimulus playbacks, suggesting that the pre-exposure led to some degree of generalization to sounds resembling pain squeaks in their frequency composition. In the second experiment, we explored whether the pre-exposure to footshocks, which triggers both an aversive inner state highly effective for fear-conditioning and the emission of pain squeaks, and is thus ideal for Hebbian learning and auto-conditioning, could be substituted using a weaker but still painful stimulus (a CO_2_ laser) paired with squeak playback. We thus employed four experimental groups, in which animals were confronted, during pre-exposure, either with (1) a painful laser stimulation paired with a squeak, (2) only the painful laser, (3) only the squeak or (4) neither stimuli. During pre-exposure, we observed reliable pain responses in the experimental groups receiving high intensity laser stimulation while these were largely absent from the other conditions. Fear responses upon the playback of pain squeaks 48h later however were much lower in case of freezing or absent in case of 22 kHz calls, with both occurring at significantly lower levels than when animals were pre-exposed to footshocks in experiment 1. There was strong evidence against a difference between the experimental groups of experiment 2 during squeak playbacks, preventing any further conclusions with respect to the auto-conditioning hypothesis.

Squeaking as a response to painful electroshocks has been documented for decades in rodents (Jourdan et al., 1995) and have recently been mechanistically investigated in mice (Ruat et al., 2022). Our study indicated that these squeaks are not innately sufficient to trigger nocifensive responses in the relatively safe environments we place the animals in, since we found no elevated rates of freezing and 22 kHz calls during squeak playback in animals without painful self-experience across both experiments. Squeaks thus have similar properties, in their dependence on prior experience, as the 22 kHz USVs (Parsana, Li, et al., 2012), or silence (Cruz et al., 2020) in other studies of emotional contagion. Since the animals in the first experiment did not only show elevated freezing during squeak but also phase-scrambled squeak replay, it is possible that the animals also became either threat-sensitized to novel stimuli in general, or generalized to the phase-scrambled squeaks given that they still shared several features of the original squeak. It is also possible that the animals underwent a form of “pseudo-conditioning” (Bouton, 2007), which results in conditioned responses in the presence of a CS that was not paired with the US.

The other proxy of fear in the animals, the 22kHz ultrasonic vocalization showed a more nuanced picture, with higher levels of ultrasonic vocalizations emitted upon hearing the intact than the phase-scrambled calls. This highlights the importance of extending our measurements of fear beyond a single behavior (freezing) to better interrogate the internal state of the animals and suggests that rats could indeed have associated somewhat specifically their own squeaks with the shock leading to a conditioned fear recall upon playback of the intact squeaks. It should be noted that there is a possibility that these somewhat specific responses are still due to sensitization as outlined by Parsana et al. (2012): responding to the squeaks could be innate and genetically hard-wired, but freezing and ultrasonic vocalizations may require the animal to have other reasons to be alert and risk aware, for this inborn sensitivity to trigger these observable behaviors – reasons the pre-exposure may have provided.

While 22 kHz vocalizations have been demonstrated to occur during solitude when exposed to predators (Blanchard et al., 1992), during fear learning and subsequent fear recall (Borta et al., 2006; Schwarting et al., 2007) or when for example subjected to aversive handling procedures (Brudzynski & Ociepa, 1992), the difference between freezing and 22 kHz calls in our results may speak to the communicative role of the 22 kHz vocalizations (Schwarting & Wöhr, 2012). Although the experiments did not feature another conspecific, the playback of squeaks and the olfactory presence of the bedding smell from other conspecifics likely induced an impression of another rat’s presence in the experimental animals since playback took place in the dark. Previous studies have demonstrated that these vocalizations can be specific alarm calls directed at conspecifics to signal potential danger in the environment (Wöhr & Schwarting, 2007). It could be speculated that the specificity of the effect could relate to freezing being more prone to sensitization as it is a self-directed response to danger whereas the 22 kHz vocalizations are less prone to sensitization as they are primarily directed towards others. Possibly the intact squeaks may have provided listeners with more reasons to communicate with a conspecific, the presence of which is suggested by the calls, than the phase scrambled squeaks. That is to say, the scrambled squeaks may still be alerting, and trigger freezing, but by being less suggestive of the presence of a conspecific, they may trigger less incentives to emit conspecific directed alarm calls. Future research is however needed to further explore this possibility.

Although the second experiment had the specific aim to shed light onto the nature of the process that generated the results from our first experiment, the results remain unfortunately inconclusive. Several possibilities come to mind as to why we did not find measurable differences between the groups that were pre-exposed to laser stimulation compared to the controls. In the study of Cruz et al. (2020), it was noted that effects of emotional contagion could only be observed if there was an aversive experience in combination with freezing behavior. Neither the aversive experience nor the freezing by itself were sufficient to induce later emotional contagion upon hearing an interruption of motion sounds. In contrast to our footshock pre-exposure, which triggers robust freezing, the laser stimulation did not produce such freezing, and may thus have triggered a state of pain without robust fear. If the animal were to have associated the pain squeaks with this state of pain without fear, the later playback of squeaks would not have triggered freezing or ultrasonic vocalization. Accordingly, our efforts to use a different stimulus (CO_2_ laser) that doesn’t trigger pain squeaks, in order to restrict auto-conditioning to the condition in which we artificially pair the pain with squeaks, appears to have backfired, because the CO_2_ laser failed to create the defensive inner state that manifests in freezing and ultrasonic vocalizations. Another possibility may be that auto-conditioning requires a self-production of the squeak. In our paradigm, the rats only listened to the playback of a squeak during pre-exposure. While neuronal activity in auditory areas might have been similar, there would have been a lack of motor expression in the laryngeal muscle that is necessary for vocalization (Riede, 2011). Thus, corresponding motor areas were not activated during laser pre-exposure. Since affective states are strongly embodied (Keysers et al., 2010; Oosterwijk et al., 2010), the mere perception of the squeak in the absence of any embodiment could have potentially impaired any conditioning to the squeak. While the results from the second experiment cannot adjudicate directly on the auto-conditioning hypothesis, they still tentatively speak against sensitization as the animals received an aversive painful experience that did not lead to any increase in freezing or 22 kHz calls. Since sensitization effects can also not explain the results on lesioning the auditory thalamus (Kim et al., 2010), we believe that an interpretation of auto-conditioning to squeaks is more likely to account for our obtained results in experiment 1.

The present study is subject to limitations that need to be acknowledged. First, the sample sizes in the first experiment are on the low side especially for the experimental group that did not receive any shocks during pre-exposure. Given the consistency of the data across animals and the support for the effect using Bayes factors, we believe that these results are valid nevertheless. Another limitation stems from the fact that the laser stimulation did not elicit fear responses that are common for shock delivery. A key difference of our laser stimulation was that its pain was much more focal than the whole-body experience resulting from footshocks. Future studies could use a wider beam or beam splitters to provide a less localized pain sensation or more frequent and intensive laser stimulations below squeak threshold. Furthermore, the close to zero latency of the squeak playback during pre-exposure might not have been optimal as heat pain might take a bit of time to actually manifest. Finally, this study was conducted exclusively in male rats. Thus, our results do not necessarily generalize to female rats. Furthermore, female rats show a different behavioral response to fearful stimuli compared to the freezing of male, i.e. darting (Gruene et al., 2015; Han et al., 2020) but also in their USV emissions (Willadsen et al., 2021). This darting behavior is constituted by brief and high velocity movements in the experimental chamber. It could be that differences that were not observed in freezing responses during squeak playback could be detected when analyzing darting behavior.

In conclusion, we could show that pre-exposure to footshocks triggers an increase in emotional contagion to hearing pain squeaks, in line with similar findings for the sound of freezing (Cruz et al., 2020) and 22 kHz calls (Parsana, Moran, et al., 2012). Indeed, we found the playback of squeaks suffices to trigger freezing levels in shock-preexposed animals that were only 20% lower than those triggered by the full experience of witnessing an animal receive the same number of footshocks in similar experiments (Han et al., 2019). We were however unable to find evidence that the pairing of pain and hearing squeaks suffices to replicate this potentiation, as auto-conditioning may have suggested. Since there seem to be multiple stimuli that rats can potentially auto-condition to, the well documented effect of prior experience on the multimodal experience of witnessing other animals in distress in close physical proximity (Keysers et al., 2022), is likely to result from cumulative effects on individual cues. This could explain why the effects of self-experience is more robust in the multimodal real-life situation (Atsak et al., 2011; Han et al., 2019; Kim et al., 2010) than when individual cues are isolated (Calub et al., 2018). In the future, our results could be complemented by for example temporally deafening the animals during shock pre-exposure using pharmacological injections. If the auto-conditioning hypothesis holds true, squeak playback should not induce fear under these conditions. Furthermore, it would be interesting to investigate the neurobiological difference between animals with and without self-experience during squeaking in emotional contagion either by playback or using a demonstrator behind an opaque divider. Here, areas such as the insula or ACC would be of particular interest due to their known contribution to emotional contagion and pain mirror responses in humans (Jabbi & Keysers, 2008; Nummenmaa et al., 2008; Soyman et al., 2021).

## Data and code availability

All data and code are accessible for review purposes and will be made fully available upon acceptance of the manuscript under the following link: https://osf.io/efuq4/.

## Author contributions

Conceptualization: EP, ES, VG, CK

Methodology: EP, ES, VG, CK

Software: JP, EP, ES

Formal Analysis: JP, EP, ES, VG, CK

Investigation: JP, ES, EP, ER, NS, SM

Resources: ES, MW

Data Curation: JP, ES, VG, CK

Writing - Original Draft: JP

Writing - Review and Editing: EP, ES, MW, VG, CK

Visualization: JP, VG, CK

Supervision: VG, CK

Project Administration: VG, CK

Funding Acquisition: VG, CK

## References

Allsop, S. A., Wichmann, R., Mills, F., Burgos-Robles, A., Chang, C.-J., Felix-Ortiz, A. C., Vienne, A., Beyeler, A., Izadmehr, E. M., Glober, G., Cum, M. I., Stergiadou, J., Anandalingam, K. K., Farris, K., Namburi, P., Leppla, C. A., Weddington, J. C., Nieh, E. H., Smith, A. C., … Tye, K. M. (2018). Corticoamygdala Transfer of Socially Derived Information Gates Observational Learning. Cell, 173(6), 1329-1342.e18. https://doi.org/10.1016/j.cell.2018.04.004

Atsak, P., Orre, M., Bakker, P., Cerliani, L., Roozendaal, B., Gazzola, V., Moita, M., & Keysers, C. (2011). Experience modulates vicarious freezing in rats: A model for empathy. PLOS ONE, 6(7), e21855. https://doi.org/10.1371/journal.pone.0021855

Baron-Cohen, S., & Wheelwright, S. (2004). The empathy quotient: An investigation of adults with Asperger syndrome or high functioning autism, and normal sex differences. Journal of Autism and Developmental Disorders, 34(2), 163–175. https://doi.org/10.1023/B:JADD.0000022607.19833.00

Blanchard, R. J., Agullana, R., McGee, L., Weiss, S., & Blanchard, D. C. (1992). Sex differences in the incidence and sonographic characteristics of antipredator ultrasonic cries in the laboratory rat (Rattus norvegicus). Journal of Comparative Psychology, 106(3), 270–277. https://doi.org/10.1037/0735-7036.106.3.270

Borta, A., Wöhr, M., & Schwarting, R. K. W. (2006). Rat ultrasonic vocalization in aversively motivated situations and the role of individual differences in anxiety-related behavior. Behavioural Brain Research, 166(2), 271–280. https://doi.org/10.1016/j.bbr.2005.08.009

Bouton, M. E. (2007). Learning and behavior: A contemporary synthesis. Sinauer Associates.

Brudzynski, S. M., Bihari, F., Ociepa, D., & Fu, X.-W. (1993). Analysis of 22 kHz ultrasonic vocalization in laboratory rats: Long and short calls. Physiology & behavior, 54(2), 215–221.

Brudzynski, S. M., & Ociepa, D. (1992). Ultrasonic vocalization of laboratory rats in response to handling and touch. Physiology & Behavior, 52(4), 655–660. https://doi.org/10.1016/0031-9384(92)90393-g

Calub, C. A., Furtak, S. C., & Brown, T. H. (2018). Revisiting the auto-conditioning hypothesis for acquired reactivity to ultrasonic alarm calls. Physiology & behavior, 194, 380–386. https://doi.org/10.1016/j.physbeh.2018.06.029

Carrillo, M., Han, Y., Migliorati, F., Liu, M., Gazzola, V., & Keysers, C. (2019). Emotional mirror neurons in the rat’s anterior cingulate cortex. Current biology, 29(8), 1301-1312. e6.

Carrillo, M., Migliorati, F., Bruls, R., Han, Y., Heinemans, M., Pruis, I., Gazzola, V., & Keysers, C. (2015). Repeated Witnessing of Conspecifics in Pain: Effects on Emotional Contagion. PLOS ONE, 10(9), e0136979. https://doi.org/10.1371/journal.pone.0136979

Choi, J.-S., & Brown, T. H. (2003). Central amygdala lesions block ultrasonic vocalization and freezing as conditional but not unconditional responses. Journal of Neuroscience, 23(25), 8713–8721.

Coffey, K. R., Marx, R. G., & Neumaier, J. F. (2019). DeepSqueak: A deep learning-based system for detection and analysis of ultrasonic vocalizations. Neuropsychopharmacology, 44(5), 859– 868.

Cruz, A., Heinemans, M., Marquez, C., & Moita, M. A. (2020). Freezing displayed by others is a learned cue of danger resulting from co-experiencing own freezing and shock. Current biology, 30(6), 1128-1135. e6.

Decety, J., & Ickes, W. (2011). The Social Neuroscience of Empathy. MIT Press.

Endres, T., Widmann, K., & Fendt, M. (2007). Are rats predisposed to learn 22 kHz calls as danger-predicting signals? Behavioural Brain Research, 185(2), 69–75.

Fendt, M., Brosch, M., Wernecke, K. E. A., Willadsen, M., & Wöhr, M. (2018). Predator odour but not TMT induces 22-kHz ultrasonic vocalizations in rats that lead to defensive behaviours in conspecifics upon replay. Scientific Reports, 8(1), 11041. https://doi.org/10.1038/s41598-018-28927-4

Friard, O., & Gamba, M. (2016). BORIS!]: a free, versatile open–source event–logging software for video/audio coding and live observations. Methods in Ecology and Evolution, 7(11), 1325– 1330. https://doi.org/10.1111/2041-210X.12584

Gruene, T. M., Flick, K., Stefano, A., Shea, S. D., & Shansky, R. M. (2015). Sexually divergent expression of active and passive conditioned fear responses in rats. Elife, 4, e11352.

Han, Y., Bruls, R., Soyman, E., Thomas, R. M., Pentaraki, V., Jelinek, N., Heinemans, M., Bassez, I., Verschooren, S., & Pruis, I. (2019). Bidirectional cingulate-dependent danger information transfer across rats. PLoS biology, 17(12), e3000524.

Han, Y., Sichterman, B., Maria, C., Gazzola, V., & Keysers, C. (2020). Similar levels of emotional contagion in male and female rats. Scientific reports, 10(1), 2763. https://doi.org/10.1038/s41598-020-59680-2

Hatfield, E., Cacioppo, J. T., & Rapson, R. L. (1993). Emotional Contagion. Current Directions in Psychological Science, 2(3), 96–100. https://doi.org/10.1111/1467-8721.ep10770953

Jabbi, M., & Keysers, C. (2008). Inferior frontal gyrus activity triggers anterior insula response to emotional facial expressions. Emotion, 8(6), 775.

Jeon, D., Kim, S., Chetana, M., Jo, D., Ruley, H. E., Lin, S.-Y., Rabah, D., Kinet, J.-P., & Shin, H.-S. (2010). Observational fear learning involves affective pain system and Cav1.2 Ca2+ channels in ACC. Nature neuroscience, 13(4), 482–488. https://doi.org/10.1038/nn.2504

Jourdan, D., Ardid, D., Chapuy, E., Eschalier, A., & Le Bars, D. (1995). Audible and ultrasonic vocalization elicited by single electrical nociceptive stimuli to the tail in the rat. PAIN\circledR, 63(2), 237–249.

Keppel, G., & Zedeck, S. (1989). Data analysis for research designs. W. H. Freeman.

Keum, S., Kim, A., Shin, J. J., Kim, J.-H., Park, J., & Shin, H.-S. (2018). A Missense Variant at the Nrxn3 Locus Enhances Empathy Fear in the Mouse. Neuron, 98(3), 588-601.e5. https://doi.org/10.1016/j.neuron.2018.03.041

Keysers, C., & Gazzola, V. (2014a). Dissociating the ability and propensity for empathy. Trends in cognitive sciences, 18(4), 163–166. https://doi.org/10.1016/j.tics.2013.12.011

Keysers, C., & Gazzola, V. (2014b). Hebbian learning and predictive mirror neurons for actions, sensations and emotions. Philosophical Transactions of the Royal Society B: Biological Sciences, 369(1644), 20130175. https://doi.org/10.1098/rstb.2013.0175

Keysers, C., Kaas, J. H., & Gazzola, V. (2010). Somatosensation in social perception. Nature Reviews Neuroscience, 11(6), 417–428. https://doi.org/10.1038/nrn2833

Keysers, C., Knapska, E., Moita, M. A., & Gazzola, V. (2022). Emotional Contagion and Prosocial Behavior in Rodents. Trends In Cognitive Sciences.

Keysers, C., Perrett, D. I., & Gazzola, V. (2014). Hebbian Learning is about contingency not contiguity and explains the emergence of predictive mirror neurons. The Behavioral and brain sciences, 37(2), 205–206. https://doi.org/10.1017/S0140525X13002343

Kim, E. J., Kim, E. S., Covey, E., & Kim, J. J. (2010). Social transmission of fear in rats: The role of 22-kHz ultrasonic distress vocalization. PLOS ONE, 5(12), e15077.

Kung, J.-C., Su, N.-M., Fan, R.-J., Chai, S.-C., & Shyu, B.-C. (2003). Contribution of the anterior cingulate cortex to laser-pain conditioning in rats. Brain research, 970(1–2), 58–72.

Lee, M. D., & Wagenmakers, E.-J. (2014). Bayesian cognitive modeling: A practical course. Cambridge university press.

Nummenmaa, L., Hirvonen, J., Parkkola, R., & Hietanen, J. K. (2008). Is emotional contagion special? An fMRI study on neural systems for affective and cognitive empathy. Neuroimage, 43(3), 571–580.

Oosterwijk, S., Topper, M., Rotteveel, M., & Fischer, A. H. (2010). When the mind forms fear: Embodied fear knowledge potentiates bodily reactions to fearful stimuli. Social Psychological and Personality Science, 1(1), 65–72.

Paradiso, E., Gazzola, V., & Keysers, C. (2021). Neural mechanisms necessary for empathy-related phenomena across species. Current Opinion in Neurobiology, 68, 107–115. https://doi.org/10.1016/j.conb.2021.02.005

Parsana, A. J., Li, N., & Brown, T. H. (2012). Positive and negative ultrasonic social signals elicit opposing firing patterns in rat amygdala. Behavioural Brain Research, 226(1), 77–86.

Parsana, A. J., Moran, E. E., & Brown, T. H. (2012). Rats learn to freeze to 22-kHz ultrasonic vocalizations through auto-conditioning. Behavioural Brain Research, 232(2), 395–399. https://doi.org/10.1016/j.bbr.2012.03.031

Pereira, A. G., Cruz, A., Lima, S. Q., & Moita, M. A. (2012). Silence resulting from the cessation of movement signals danger. Current biology, 22(16), R627–R628.

Pérez-Manrique, A., & Gomila, A. (2022). Emotional contagion in nonhuman animals: A review. Wiley interdisciplinary reviews. Cognitive science, 13(1), e1560. https://doi.org/10.1002/wcs.1560

Poulos, A. M., Zhuravka, I., Long, V., Gannam, C., & Fanselow, M. (2015). Sensitization of fear learning to mild unconditional stimuli in male and female rats. Behavioral neuroscience, 129(1), 62.

Riede, T. (2011). Subglottal pressure, tracheal airflow, and intrinsic laryngeal muscle activity during rat ultrasound vocalization. Journal of neurophysiology, 106(5), 2580–2592.

Ruat, J., Genewsky, A. J., Heinz, D. E., Kaltwasser, S. F., Canteras, N. S., Czsich, M., Chen, A., & Wotjak, C. T. (2021). Why do mice squeak? Towards a better understanding of defensive vocalization. iScience.

Schwarting, R. K. W., Jegan, N., & Wöhr, M. (2007). Situational factors, conditions and individual variables which can determine ultrasonic vocalizations in male adult Wistar rats. Behavioural Brain Research, 182(2), 208–222. https://doi.org/10.1016/j.bbr.2007.01.029

Schwarting, R. K. W., & Wöhr, M. (2012). On the relationships between ultrasonic calling and anxiety-related behavior in rats. Brazilian Journal of Medical and Biological Research, 45(4), 337–348. https://doi.org/10.1590/S0100-879X2012007500038

Shyu, B.-C., Chai, S.-C., Kung, J.-C., & Fan, R.-J. (2003). A quantitative method for assessing of the affective component of the pain: Conditioned response associated with CO2 laser-induced nocifensive reaction. Brain research protocols, 12(1), 1–9.

Soyman, E., Bruls, R., Ioumpa, K., Müller-Pinzler, L., Gallo, S., van Straaten, E. C. W., Self, M. W., Peters, J. C., Possel, J. K., & Onuki, Y. (2021). Intracortical human recordings reveal intensity coding for the pain of others in the insula. bioRxiv.

Terranova, J. I., Yokose, J., Osanai, H., Marks, W. D., Yamamoto, J., Ogawa, S. K., & Kitamura, T. (2022). Hippocampal-amygdala memory circuits govern experience-dependent observational fear. Neuron, 110(8), 1416-1431.e13. https://doi.org/10.1016/j.neuron.2022.01.019

Willadsen, M., Üngör, M., Sługocka, A., Schwarting, R. K. W., Homberg, J. R., & Wöhr, M. (2021). Fear Extinction and Predictive Trait-Like Inter-Individual Differences in Rats Lacking the Serotonin Transporter. International Journal of Molecular Sciences, 22(13), 7088. https://doi.org/10.3390/ijms22137088

Wöhr, M., & Schwarting, R. K. W. (2007). Ultrasonic communication in rats: Can playback of 50-kHz calls induce approach behavior? PloS One, 2(12), e1365. https://doi.org/10.1371/journal.pone.0001365

Wöhr, M., & Schwarting, R. K. W. (2008). Ultrasonic calling during fear conditioning in the rat: No evidence for an audience effect. Animal Behaviour, 76(3), 749–760. https://doi.org/10.1016/j.anbehav.2008.04.017

