## Supplementary Information for "Investigating the mechanistic role of painful self-experience in emotional contagion: an effect of auto-conditioning?"

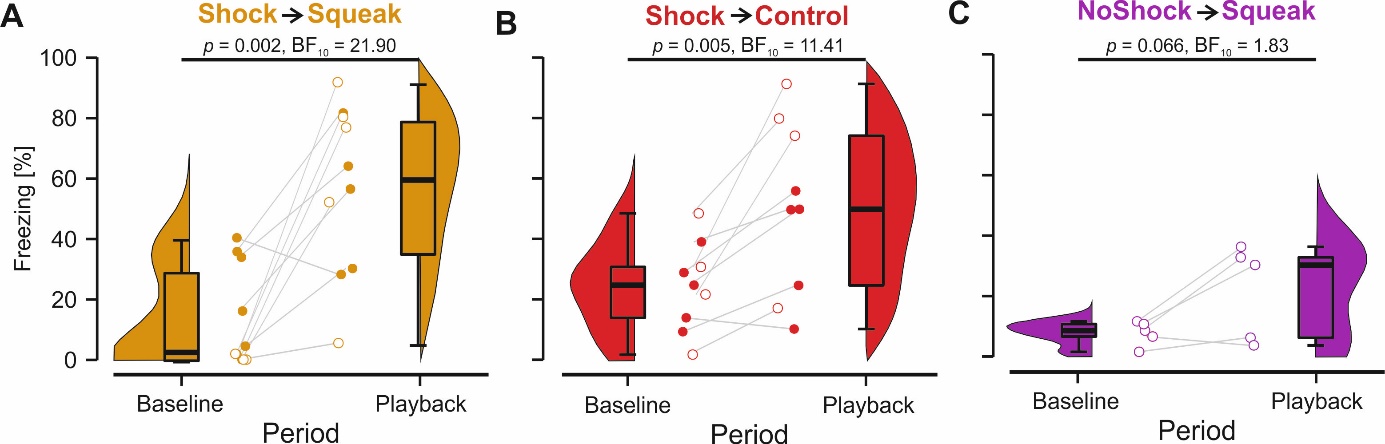

**Supplementary Figure 1.** Proportion of freezing responses in percent during baseline and auditory playback in the playback session. Paired tests (t-test, if the Shapiro-Wilk (S-W) normality test did not reject normality, i.e. S-W p-value>0.05 or Wilcoxon signed-rank, if it did, i.e. S-W p<0.05) indicated that both the **A**) Shock->Squeak group (S-W p-value = 0.773; t(9) = 4.26, p = 0.002, BF_10_ = 21.90) and the **B**) Shock->Control group (S-W p-value = 0.666; t(8) = 3.86, p = 0.005, BF_10_ 11.41) increased their freezing from baseline to the playback period whereas the **C**) NoShock->Squeak group showed a trend towards a significant increase (S-W p-value = 0.207; t(4) = 2.52, p = 0.066, BF_10_ = 1.83). Animals with higher amplitude playback are marked by an open circle. The p-values indicated for the changes from baseline to playback were not corrected for multiple comparisons, but applying an FDR correction would not alter the conclusion.

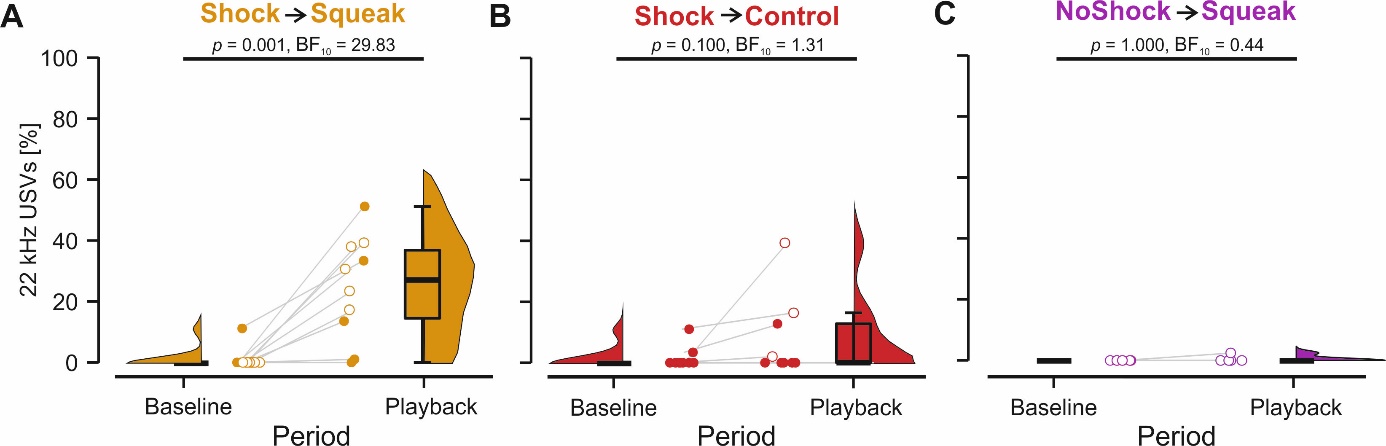

**Supplementary Figure 2.** Proportion of 22 kHz calls in percent during baseline and auditory playback in the playback session. For 22 kHz calls, only the **A**) Shock->Squeak group showed an increase from baseline to playback (S-W p-value = 0.852; t(9) = 4.51, p = 0.001, BF_10_ = 29.83). The **B**) Shock->Control (S-W p-value < 0.001; z = 1.83, p = 0.100, BF_10_ = 1.31) and **C**) NoShock->Squeak (S-W p-value < 0.001; z = 0.48, p = 1.000, BF_10_ = 0.44) showed no significant change. Animals with higher amplitude playback are marked by an open circle. The p-values indicated for the changes from baseline to playback were not corrected for multiple comparisons, but applying an FDR correction would not alter the conclusion

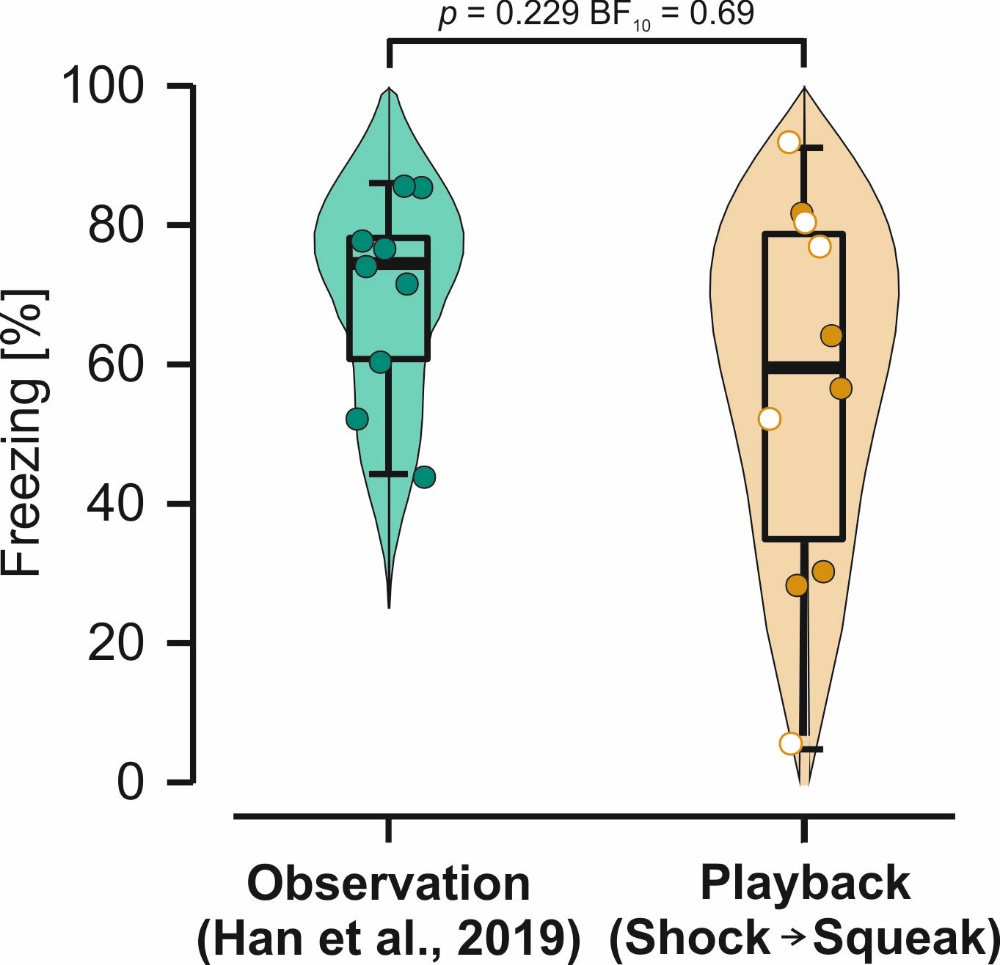

**Supplementary Figure 3.** Proportion of freezing responses comparing the results from Han et al. (2019) that used a real-life demonstrator during emotional contagion vs. our Shock -> Squeak group that merely played back the squeak.

.

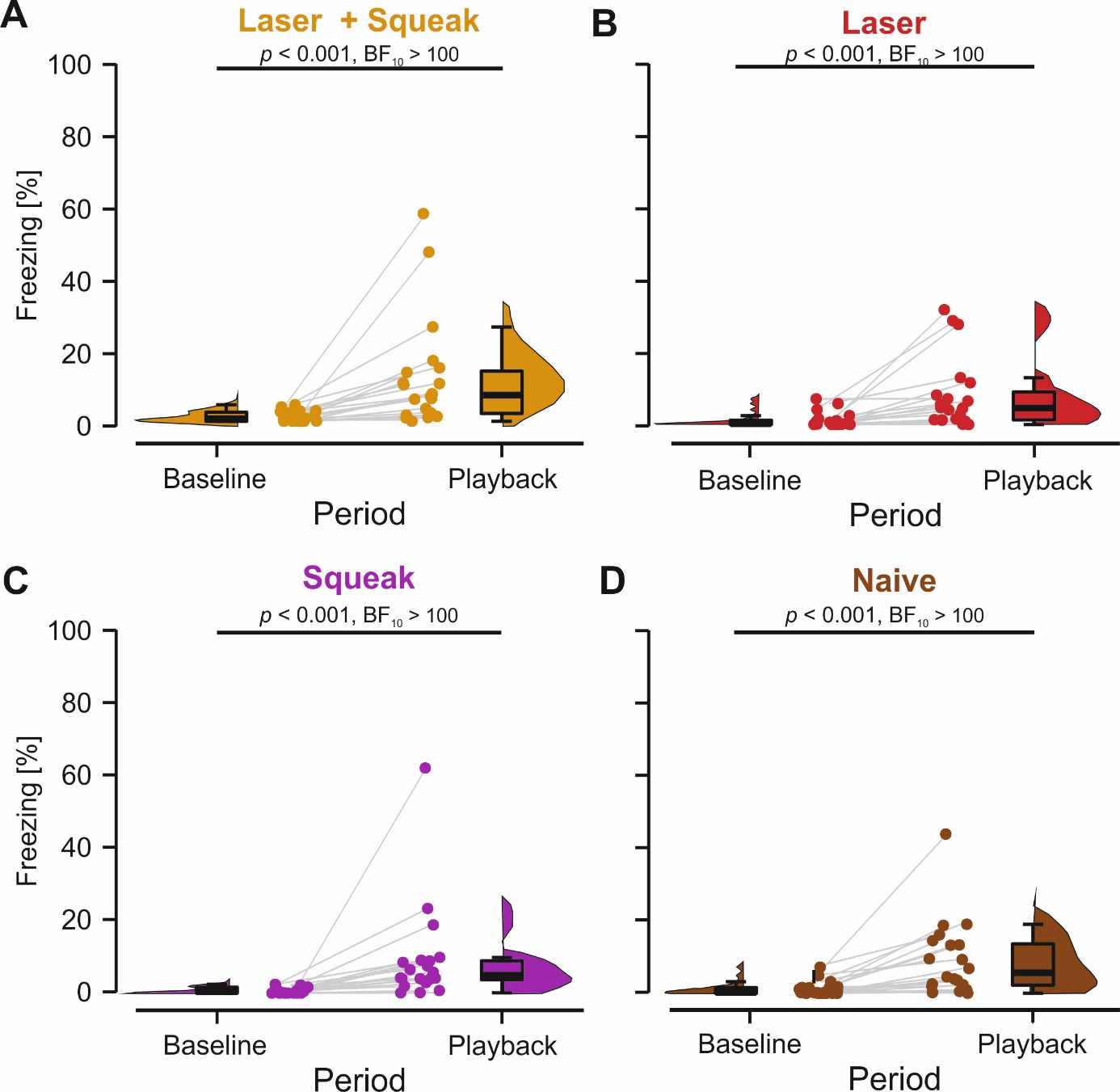

**Supplementary Figure 4.** Proportion of freezing responses in percent during the squeak playback session. Paired Wilcoxon signed-rank tests indicated extreme evidence for the **A**) Laser + Squeak group (S-W p-value < 0.001; z = 3.82, p < 0.001, BF_10_ > 100), the **B**) Laser group S-W p-value < 0.001; z = 3.81, p < 0.001, BF_10_ > 100), the **C**) Squeak group S-W p-value < 0.001; z = 3.92, p < 0.001, BF_10_ > 100), and the **D**) Naïve group (S-W p-value < 0.001; z = 3.61, p < 0.001, BF_10_ > 100) for an increase in freezing from the baseline to the playback period. The p-values indicated for the changes from baseline to playback were not corrected for multiple comparisons, but applying an FDR correction would not alter the conclusion

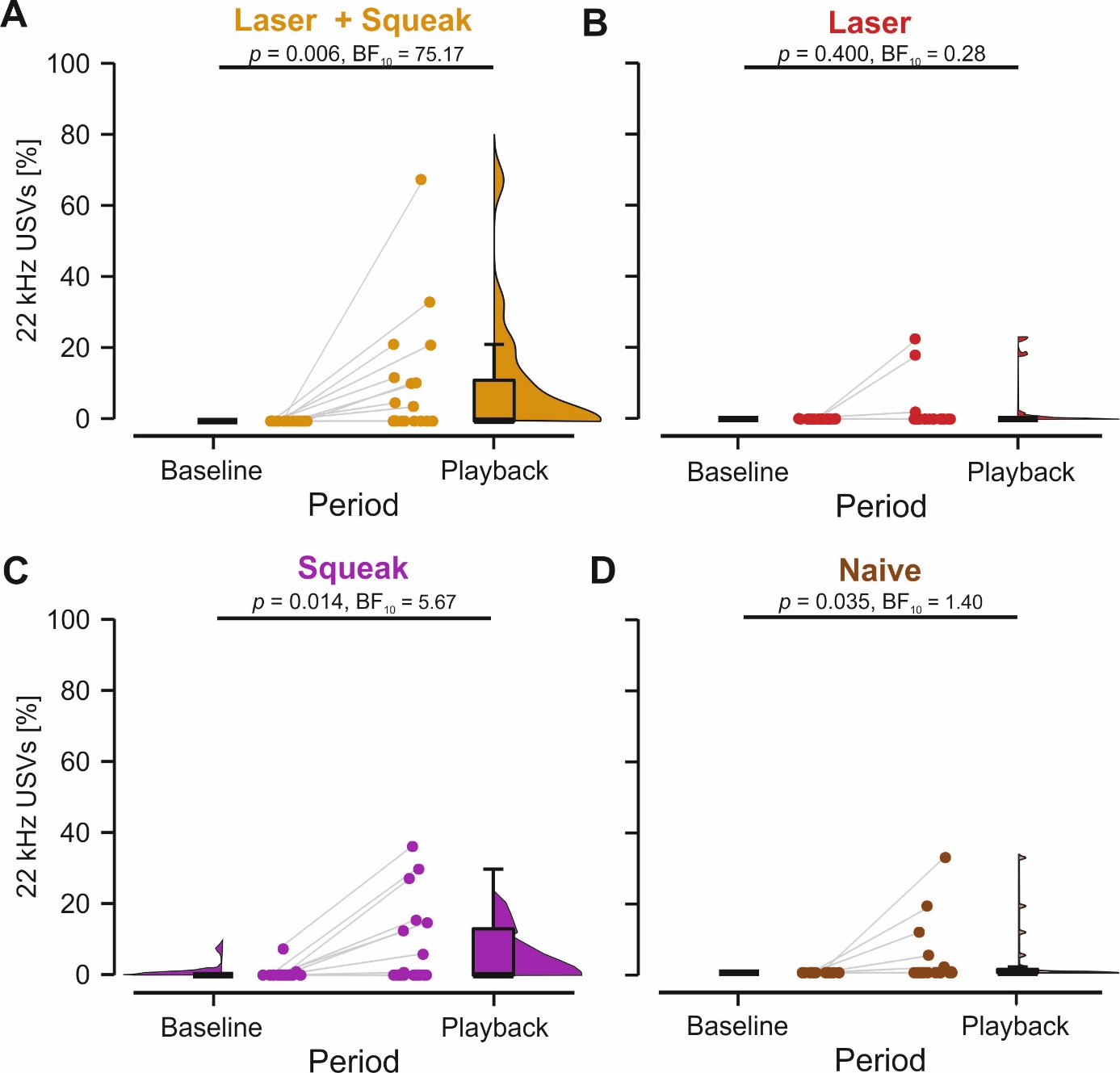

**Supplementary Figure 5.** Proportion of 22 kHz calls in percent during the squeak playback session. The **A)** Laser+Squeak group demonstrated strong to extreme evidence for an increase from the baseline to the playback period (S-W p-value < 0.001; z = 2.80, p = 0.006, BF_10_ = 75.17) whereas moderate evidence of absence of this effect was observed for the **B)** Laser group (S-W p-value < 0.001; z = 0.94, p = 0.400, BF_10_ = 0.28). For the **C)** Squeak group, we found moderate evidence in favor of an increase in 22 kHz calls (S-W p-value < 0.001; z = 2.52, p = 0.014, BF_10_ = 5.67). No evidence in either direction was found for the **D)** Naïve group (S-W p-value < 0.001; z = 2.20, p = 0.035, BF_10_ = 1.40). The p-values indicated for the changes from baseline to playback were not corrected for multiple comparisons, but applying an FDR correction would not alter the conclusion

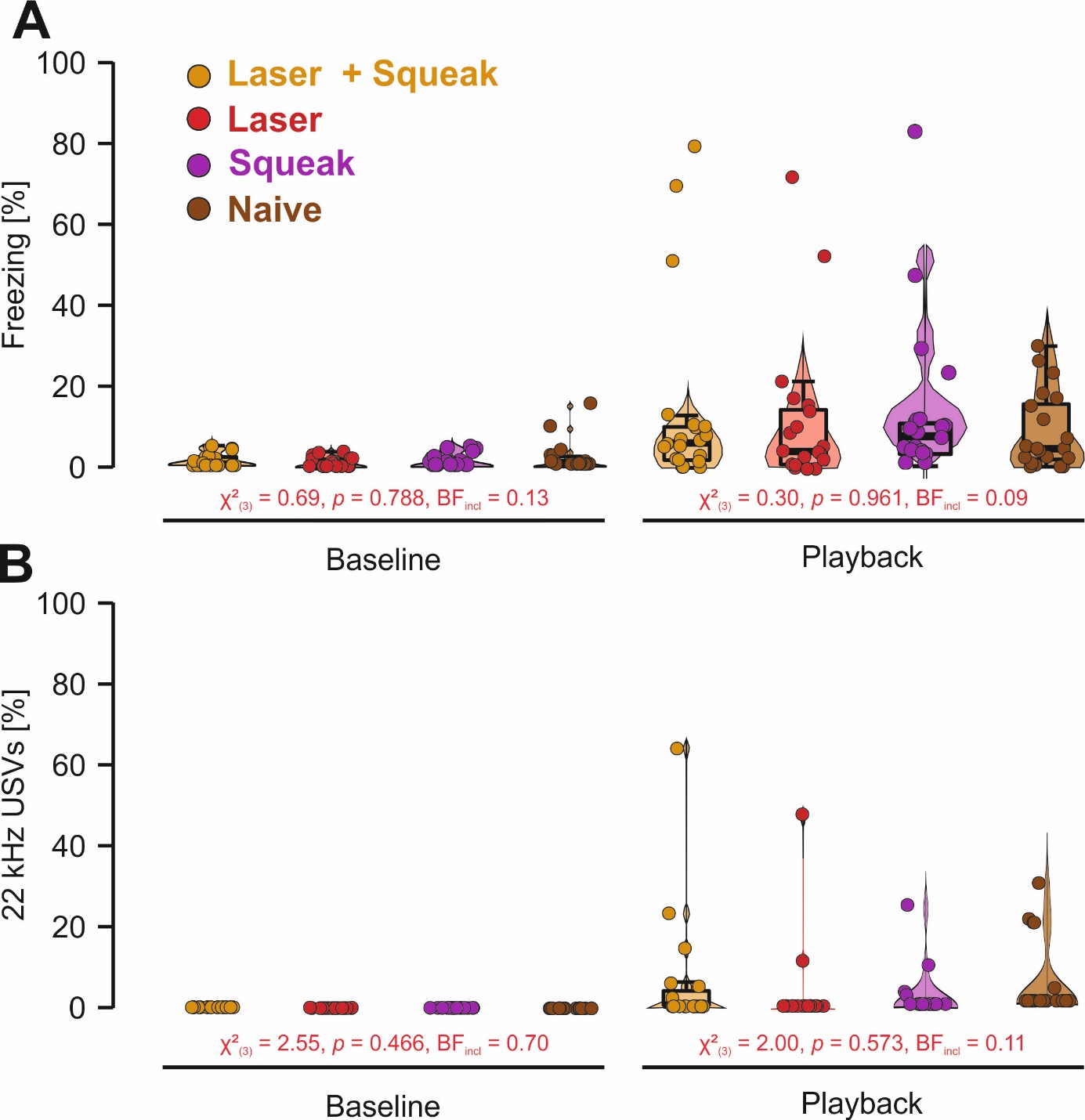

**Supplementary Figure 6. A)**  Proportion of freezing responses in percent during baseline and auditory playback in the playback session on day 5. Animals with higher amplitude playback are marked by an open circle. Only between group differences are presented here. Within group differences from baseline to playback are depicted in Supplementary Figure 7. **B)** Proportion of 22 kHz vocalizations in percent during baseline and auditory playback in the playback session on day 5. Animals with higher amplitude playback are marked by an open circle. Only between group differences are presented here. Within group differences from baseline to playback are depicted in Supplementary Figure 8.

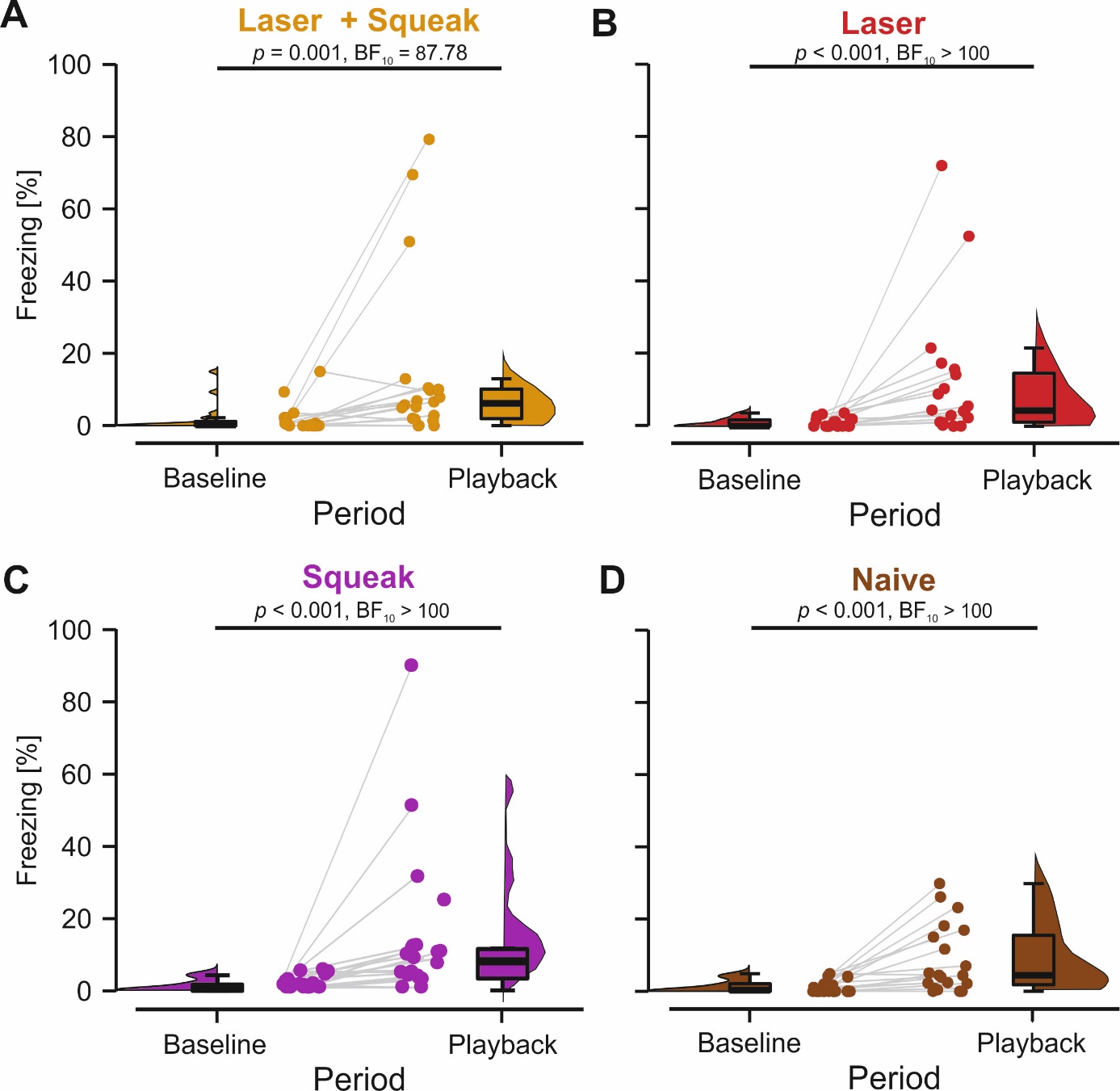

**Supplementary Figure 7.** Proportion of freezing responses in percent during the phase-scrambled squeak playback session. Paired Wilcoxon signed-ranked tests indicated strong to extreme evidence for the **A**) Laser + Squeak group (S-W p-value < 0.001; z = 3.25, p = 0.001, BF_10_ > 87.78), the **B**) Laser group S-W p-value < 0.001; z = 3.64, p < 0.001, BF_10_ > 100), the **C**) Squeak group S-W p-value < 0.001; z = 3.66, p < 0.001, BF_10_ > 100), and the **D**) Naïve group (S-W p-value < 0.001; z = 3.57, p < 0.001, BF_10_ > 100) for an increase in freezing from the baseline to the playback period. The p-values indicated for the changes from baseline to playback were not corrected for multiple comparisons, but applying an FDR correction would not alter the conclusion

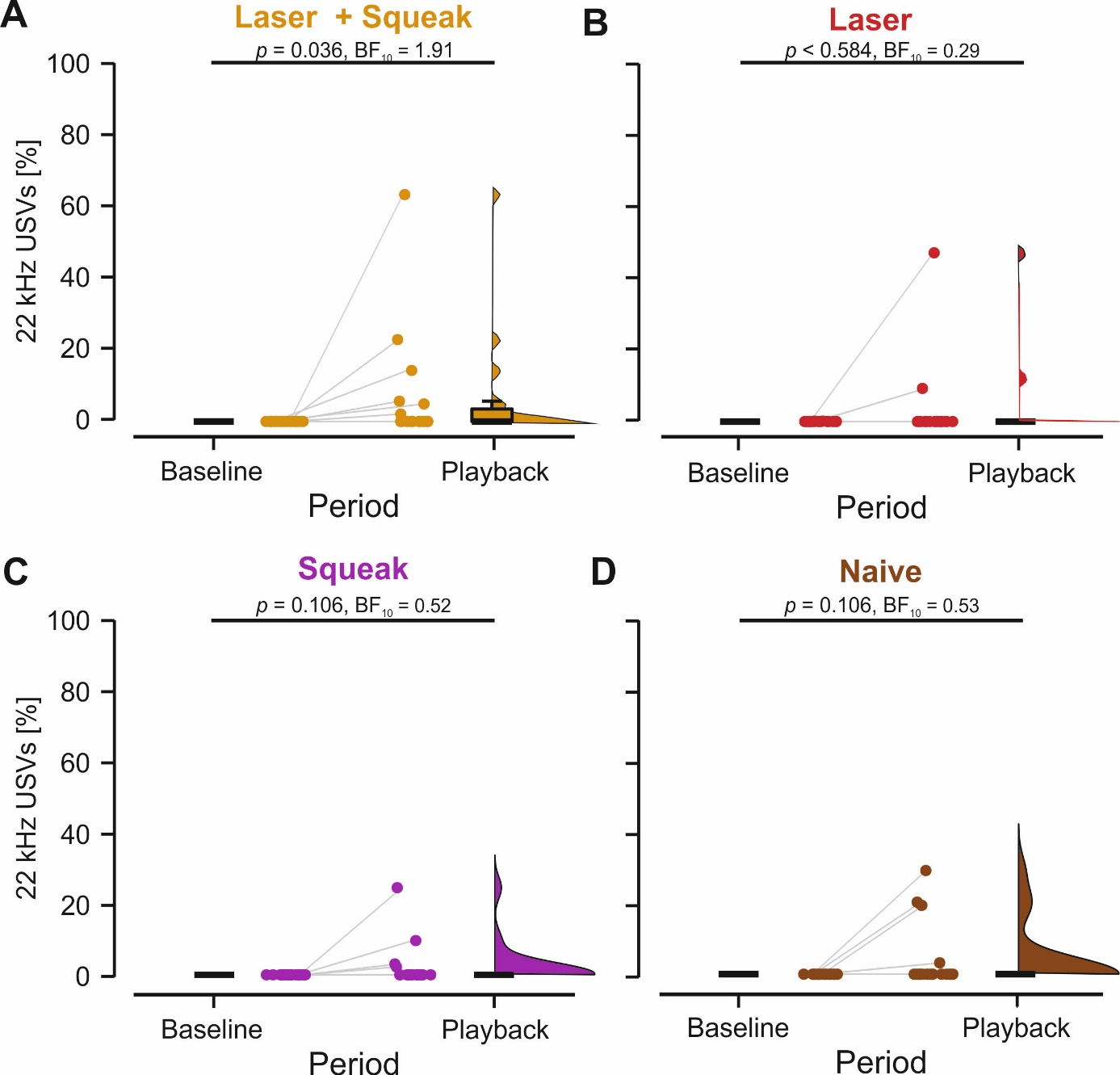

**Supplementary Figure 8.** Proportion of freezing responses in percent during the phase-scrambled squeak playback session. We found a significant increase from baseline to playback in the **A)** Laser+Squeak group (S-W p-value < 0.001; z = 2.2.20, p = 0.036).The evidence for this increase was however only anecdotal (BF_10_ = 1.91). For the **B)** Laser group there was moderate evidence of absence for an increase in 22 kHz calls during playback compared to baseline (S-W p-value < 0.001; z = 0.73, p = 0.584, BF_10_ = 0.29). For the **C)** Squeak group (S-W p-value < 0.001; z = 1.75, p = 0.106, BF_10_ = 0.52) and the **D)** Naïve group (S-W p-value < 0.001; z = 1.75, p = 0.106, BF_10_ = 0.53), there was absence of evidence for any effect. The p-values indicated for the changes from baseline to playback were not corrected for multiple comparisons, but applying an FDR correction would not alter the conclusion

**Supplementary Table 1.** Descriptive values for the proportions of freezing for the baseline and auditory playback period across the experimental groups in experiment 1.

| **Proportion of Freezing during Auditory Playback** | | | |
| --- | --- | --- | --- |
| **Trial** | **Group** | **Mean** | **SD** |
|  | Shock->Squeak | 13.4 | 16.8 |
| Baseline | Shock->Control | 24.2 | 14.6 |
|  | NoShock->Squeak | 7.8 | 4 |
|  | Shock->Squeak | 56.7 | 28 |
| Playback | Shock->Control | 50.3 | 28.6 |
|  | NoShock->Squeak | 21.9 | 15.6 |

**Supplementary Table 2.** Descriptive values for the proportions of 22 kHz calls for the baseline and auditory playback period across the experimental groups in experiment 1.

| **Proportion of 22 kHz Calls during Auditory Playback** | | | |
| --- | --- | --- | --- |
| **Trial** | **Group** | **Mean** | **SD** |
|  | Shock->Squeak | 1.1 | 3.5 |
| Baseline | Shock->Control | 1.6 | 3.7 |
|  | NoShock->Squeak | 0 | 0 |
|  | Shock->Squeak | 24.8 | 16.8 |
| Playback | Shock->Control | 7.8 | 13.4 |
|  | NoShock->Squeak | 0.5 | 1.1 |

**Supplementary Table 3.** Descriptive values for the pain reactions to the laser during pre-exposure in experiment 2.

| **Cumulative Pain Reactions during Pre-exposure** | | |
| --- | --- | --- |
| **Group** | **Mean** | **SD** |
| Laser+Squeak | 9.1 | 2 |
| Laser | 9.4 | 1.5 |
| Squeak | 2.1 | 1.9 |
| Naive | 0.9 | 1.1 |

**Supplementary Table 4.** Descriptive values for the proportions of freezing during pre-exposure in experiment 2.

| **Proportion of Freezing during Pre-exposure** | | |
| --- | --- | --- |
| **Group** | **Mean** | **SD** |
| Laser+Squeak | 24.9 | 20.4 |
| Laser | 26.8 | 21.3 |
| Squeak | 26.9 | 20.5 |
| Naive | 17 | 12.5 |

**Supplementary Table 5.** Descriptive values for the proportions of freezing during pre-exposure in experiment 2.

| **Proportion of 22 kHz Calls during Pre-exposure** | | |
| --- | --- | --- |
| **Group** | **Mean** | **SD** |
| Laser+Squeak | 0.8 | 2.6 |
| Laser | 0.4 | 1.9 |
| Squeak | 0 | 0 |
| Naive | 0.8 | 3.8 |

**Supplementary Table 6.** Descriptive values for the proportions of freezing for the baseline and auditory playback period during regular squeak playback across the experimental groups in experiment 2.

| **Proportion of Freezing during Squeak Playback** | | | |
| --- | --- | --- | --- |
| **Trial** | **Group** | **Mean** | **SD** |
|  | Laser+Squeak | 1.7 | 2 |
| Baseline | Laser | 1.3 | 2.1 |
|  | Squeak | 0.9 | 1.1 |
|  | Naive | 1.8 | 2.5 |
|  | Laser+Squeak | 15 | 17.6 |
| Playback | Laser | 8.6 | 9.6 |
|  | Squeak | 10.3 | 13.7 |
|  | Naive | 10.6 | 12.3 |

**Supplementary Table 7.** Descriptive values for the proportions of 22 kHz calls for the baseline and auditory playback period during regular squeak playback across the experimental groups in experiment 2.

| **Proportion of 22 kHz Calls during Squeak Playback** | | | |
| --- | --- | --- | --- |
| **Trial** | **Group** | **Mean** | **SD** |
|  | Laser+Squeak | 0 | 0 |
| Baseline | Laser | 0.1 | 1.4 |
|  | Squeak | 1.2 | 1.7 |
|  | Naive | 0.4 | 1.6 |
|  | Laser+Squeak | 9.9 | 17.1 |
| Playback | Laser | 2.2 | 6.4 |
|  | Squeak | 7.1 | 11.6 |
|  | Naive | 3.7 | 8.5 |

**Supplementary Table 8.** Descriptive values for the proportions of freezing for the baseline and auditory playback period during phase-scrambled squeak playback across the experimental groups in experiment 2.

| **Proportion of Freezing during Phase-Scrambled Squeak Playback** | | | |
| --- | --- | --- | --- |
| **Trial** | **Group** | **Mean** | **SD** |
|  | Laser+Squeak | 2 | 3.8 |
| Baseline | Laser | 1.1 | 1.4 |
|  | Squeak | 1.2 | 1.7 |
|  | Naive | 1.6 | 2.3 |
|  | Laser+Squeak | 15 | 23.1 |
| Playback | Laser | 13.2 | 18.7 |
|  | Squeak | 14.1 | 20.9 |
|  | Naive | 9.8 | 10.1 |

**Supplementary Table 9.** Descriptive values for the proportions of 22 kHz calls for the baseline and auditory playback period during phase-scrambled squeak playback across the experimental groups in experiment 2.

| **Proportion of 22 kHz Calls during Phase-Scrambled Squeak Playback** | | | |
| --- | --- | --- | --- |
| **Trial** | **Group** | **Mean** | **SD** |
|  | Laser+Squeak | 0 | 0 |
| Baseline | Laser | 0.1 | 0.4 |
|  | Squeak | 0 | 0 |
|  | Naive | 0 | 0 |
|  | Laser+Squeak | 6 | 15.2 |
| Playback | Laser | 3.2 | 11.3 |
|  | Squeak | 2.2 | 6 |
|  | Naive | 3.8 | 8.7 |
